# Growth, organic matter release, aggregation and recycling during a diatom bloom: A model-based analysis of a mesocosm experiment

**DOI:** 10.1101/2022.05.18.492269

**Authors:** Onur Kerimoglu, Nils H. Hintz, Leonhard Lücken, Bernd Blasius, Lea Böttcher, Carina Bunse, Thorsten Dittmar, Benedikt Heyerhoff, Corinna Mori, Maren Striebel, Meinhard Simon

## Abstract

Mechanisms terminating phytoplankton blooms are often not well understood. Potentially involved processes such as consumption by grazers, flocculation, and viral lysis each have different post-bloom consequences on the processing of the organic material, therefore it is important to develop a better understanding of the relevance of these processes, and potential interactions between them. In this study, we present a model-based analysis of a spring bloom observed in a mesocosm experiment. The intermediate-complexity (27-state variable) numerical model we extended from an earlier version to this end can resolve C, N, P and Si cycles, and relevant processes like formation of various organic material size classes (low and high molecular weight (hereafter small and large) dissolved organic carbon (DOC), transparent exopolymer particles (TEP), and small/large detritus) and their degradation by two bacterial sub-communities (free-living and particle-attached) and planktonic protists (heterotrophic flagellates and ciliates). The model can explain *>* 90% of the variation of a rich set of observations consisting of 11 independent variables over the course of 13 days during which a bloom largely dominated by diatoms develops, and disappears almost entirely. Fluxes estimated by the model point to the importance of coagulation (TEP formation) as a sink term for DOC, and a source term for POC. Consequently, aggregation with TEPs constitute an important loss term for phytoplankton. The flocculated phytoplankton, and detrital material, in turn become rapidly degraded by the particle attached bacteria and other protist heterotrophs. Through a scenario analysis, the relevance of nutrient-stress enhanced lysis rates; alterations between small and large DOC in phytoplankton exudates; and coagulation of smaller DOC molecules were investigated. Our results suggest that the former two processes have negligible effects in isolation, but when combined with the latter, they can synergistically cause substantial deviations in TEP formation, hence, in flocculation rates; and consequently in the peak magnitude of the diatom bloom, and in timing of its termination. Our results point to a need for better understanding of processes governing the termination of phytoplankton blooms, their inter-dependencies, and consequences on the global biogeochemical cycles.

## 1 Introduction

In aquatic and marine ecosystems in temperate climate zones, phytoplankton commonly form ‘blooms’ during spring. Reasons for the abrupt increase in their biomass are relatively well understood: after an onset of stratification and thereby limitation of vertical transport, phytoplankton maintained within the surface layers experience increasingly better light conditions (Huisman et al., 1999; Siegel et al., 2002). When this is combined with abundantly available dissolved inorganic nutrients, which have been building up throughout the winter (Huppert et al., 2002; Peeters et al., 2013), a so called ‘exponential growth phase’ begins. These blooms often end abruptly as well, but the reasons thereof are relatively less understood (Sommer et al., 2012), because of a multitude of factors involved, such as, but not limited to, cease of growth due to nutrient depletion, and various loss mechanisms such as the consumption by zooplankton (e.g., Irigoien et al., 2005; Wiltshire et al., 2008), flocculation (e.g., Riebesell, 1991; Passow et al., 1994), and as increasingly recognized, viral infections (e.g., Lehahn et al., 2014; Kranzler et al., 2019). Based on the Chl-*a* measurements only, which is typically the only systematically available data source apart from the nutrient concentrations, it is often not possible to identify the relative importance of these various loss processes. Such an identification is important, because these mechanisms can have substantially different consequences on the fate and recycling of the organic material later in the season.

In this study, we focused on a spring bloom observed in a mesocosm experiment. Starting from initial nutrient concentrations and phytoplankton assemblages in the coastal North Sea, and forcing the mesocosms with temperature and diurnally varying irradiance set up to reflect the natural field conditions, and given the absence of vertical heterogeneity within the tanks, a rapidly developing spring bloom was obtained within the first few days of the experiment, as reflected by the particulate organic matter (POM) and Chlorophyll-*a* (hereafter Chl-*a*) concentrations (see also Mori et al., 2021). This was followed by a sudden collapse of the phytoplankton biomass within a few days, again, as evidenced by POM and Chl-*a* concentrations.

A rich observation set was acquired throughout the experiment, both in terms of the variety and temporal resolution of measurements (Mori et al., 2021). However, based on this data set, it is still not possible to disentangle the exact mechanisms causing the collapse of the bloom. Following the depletion of nutrients, transition from growth to senescence is as expected, but the exact reasons for the rapid collapse of the diatom bloom and POM are not immediately clear. Zooplankton consumption can be ruled out as a candidate mechanism, as in the experiment under consideration, mesozooplankton were excluded by prefiltration, and other protist feeders, such as the heterotrophic flagellates and ciliates were in very low concentrations when the termination of the bloom began. Remaining candidate mechanisms include increasing lysis rates, and aggregation/flocculation processes, for which macroobservational evidence exist (Fig. SI-3).

Our main motivation here was to construct a mechanistic numerical model that can provide a plausible representation of the system, such that it can then be used to bridge the gaps between the available measurements, and allow quantifying fluxes between various components of the system, as well as exploring potential inter-dependencies between various loss processes, and the consequences thereof for the recycling of the organic material. Such a modelling approach can also help improving the general understanding of the mechanisms leading to the collapse of phytoplankton blooms in temperate seas.

## 2 Methods

### 2.1 The Mesocosm Experiment and observations

The mesocosm experiment was conducted in four 600 L steel tanks, namely the ‘planktotrons’ (Gall et al., 2017), that were frequently sampled (for some variables, at subdaily intervals). The inoculum was collected from the coastal North Sea, preincubated for 10 days and then transferred to an artificial seawater medium with a very low DOC background, taken from a nearby tidal basin of the Wadden Sea (54^*◦*^4.55’N, 7^*◦*^37.62’E). Zooplankton larger than 100*µ*m were excluded by prefiltration of the inoculum, to avoid stochastic effects of a few animals that would be otherwise present in each mesocosm. The total duration of the experiments was 38 days (March 21^st^ to April 26^th^, 2018), and included two consecutive blooms, an initial diatom bloom and after its collapse a bloom dominated greatly by *Phaeocystis globosa*. In the current study, we focus on the initial diatom bloom that took place within the first ∼15 days of the experiment. Within this period, variation between the replicates were negligable, therefore the planktotrons are not represented individually in this study, but in aggregate form (i.e., in terms of averages and standard deviations across the four planktotrons). Regarding light and temperature, field conditions were mimicked including the seasonal rise of incoming photosynthetically active radiation and water temperatures, as well as the diurnal (14:10) light cycles. The planktotrons were mixed once an hour using rotating paddles spread over the whole depth of the tanks. Details of the experimental setup, as well as the sampling procedures can be found in Mori et al. (2021). In the following, most essential information is briefly summarized.

#### 2.1.1 Abiotic and biotic variables assessed

##### Nutrient concentrations

sampling and analytical procedures for nitrate 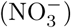, nitrite 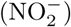, ammonium 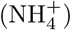, dissolved inorganic phosphorus and silicate (DIP and DISi) are described by Mori et al. (2021).

##### TDN, DOC, DOC_S_ and DOC_L_

we considered DOC and total dissolved nitrogen (TDN) measurements that were available at 18:00 each day (see Mori et al. (2021) for details). For DOC, measurements on the 23rd and 31st March at 18:00 were missing in planktotrons 2 and 4, respectively, which were filled by linear interpolation, prior to averaging the data across the planktotrons. DON was calculated by substracting total 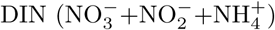 from the TDN cs. To be able to distinguish the mechanisms generating, destroying and aggregating different DOC molecules (e.g., Engel et al., 2004; Schartau et al., 2007; Polimene et al., 2018), we used a size-based approach with two classes (see also Section 2.2). The small DOC class, DOC_S_ refers to the low molecular weight (LMW)includes monomeric amino acids and monosaccharides (and other small organic molecules (*<*500 Dalton), whereas the large DOC class, DOC_L_ consists of high molecular weight (HMW) oligo- and polymeric amino acids and polysaccharides and other organic molecules (*>*500 Dalton), measurement of which are explained by Dlugosch et al. (in prep). Lipids and complex glycoproteins, which were not detected by our HPLC methods, in reality contribute also to the HMW DOC. We quantified DOC_L_, and substracting it from the DOC, we obtained the concentration of small DOC class, DOC_S_.

##### POM (C,N,P)

Sampling of POM is described by Mori et al. (2021).

##### Chl-*a*

was measured in accordance to (Thrane et al., 2015), photometrically in 96-well, F-bottom polystyrene plates (Micro plates, Boettger, Germany). Preceding pigment extraction was performed within 2 mL 90% EtOH and sonication for 30 min on ice. Further extraction took place for at least 20 h at 6^*◦*^C in the dark. Spectral absorbance was determined via plate reader (Synergy H1, BioTek instruments, VT USA) at a wavelength range of 400-800 nm in 1 nm steps. All samples were measured in technical triplicates of 200 *µ*L respectively.

##### Microscopic counts and biomass conversion of plankton

For the identification and counting of plankton, samples were conserved with Lugol’s iodine solution (1% vol.). Identification of community composition was performed via Utermöhl’s method (Utermöhl, 1931, 1958) with an inverted microscope (CKX41, Olympus Europa se & co. kg, Germany). Depending on species abundance, magnification was adjusted individually with a minimum number of 500 counted individuals per sample. Calculation of cellular biovolume is based on local time-series data provided by the Lower Saxon State Department for Waterway, Coastal and Nature Conservation (NLWKN). For rare species, literature data by (Olenina et al., 2006), the PEG Biovolume list of 2019 (http://www.ices.dk/marine-data/Documents/ENV/PEG_BVOL.zip) and North-Sea specific literature (Kraberg et al., 2010) were used. For the heterotrophic flagellates, calculation of carbon biomass was performed based on the taxanomic and allometric relationships according to Menden-Deuer and Lessard (2000). For ciliates, first species and size-class specific average biovolumes were estimated based on the available geometric information, then the carbon biomass was estimated assuming an average 0.19 pgC mm^*−*3^ (Yang et al., 2014).

##### Bacteria (free-living) counts and biomass conversion

Bacteria cell counts were measured with a flow cytometer as described in Mori et al. (2021) and Dlugosch et al (in prep.). Carbon biomass of bacteria was calculated by assuming 20 fgC cell^*−*1^ (Lee and Fuhrman, 1987).

### 2.2 The Model

We developed an ordinary differential equation model of intermediate complexity (27 state variables) and applied in zero-dimensions (0-D) to simulate the mesocosm experiment. The main components of the model is shown in Fig. 1 (see Fig. SI-4 for a representation of simulated model fluxes). A detailed description of the model is provided in Section SI-1.1 in Supporting Information.

**Figure 1.**
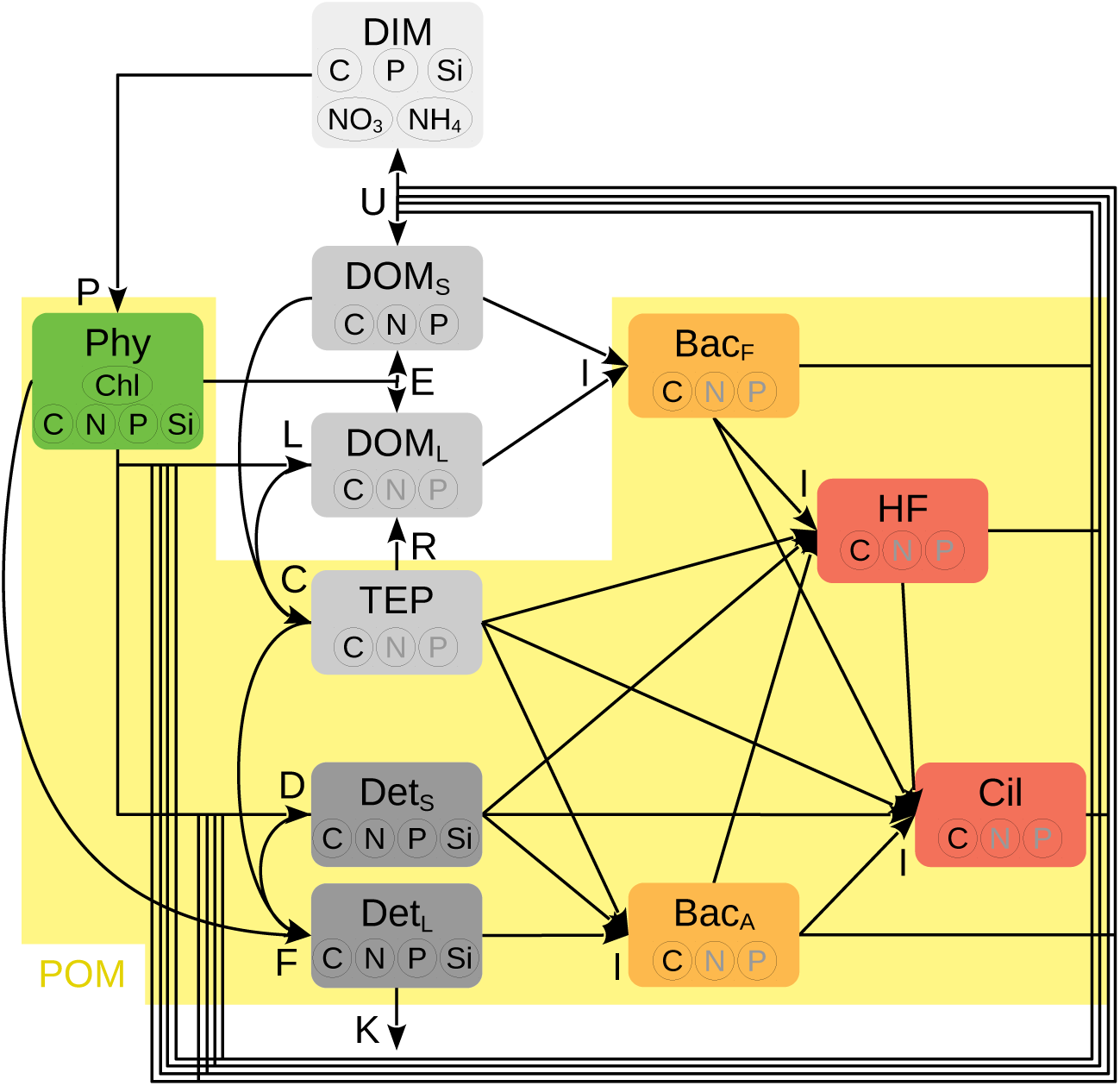
Simplified model diagram of material flows. **Abbreviations** inside boxes represent pools (POM: particulate organic Material; Phy: phytoplankton; DIM: dissolved inorganic material; DOM_S/L_: small and large dissolved organic material; TEP: transparent exopolymer particles; Det_S/L_: small and large detritus (flocs); Bac_F/A_.: free-living and particle-attached bacteria; HF: heterotrophic flagellates; Cil: ciliates) and **letters** next to arrow heads represent processes (P: production; E: exudation; L: lysis; D: death; I: ingestion; U: unassimilated ingestion; C: coalescence; R: degradation; F: flocculation; K: leakage). Biochemical content of each pool considered by the model are indicated by the inset disks, where black and gray letters represent the dynamically and diagnostically traced quantities, respectively. For details, see Section SI-1.1.

Phytoplankton, which during the spring bloom we are addressing here almost entirely consist of diatoms, take up nutrients available in dissolved inorganic form (i.e., DIC, DIP, DISi, 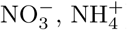) for production (P in Fig. 1), and exudate (E) carbon rich organic substances proportional to this production rate, through a process known as ‘carbon overconsumption’ (Toggweiler, 1993). These exudates consist mainly of small molecules (i.e., low molecular weight (LMW) compounds and monomers), and are therefore directed to DOM_S_ at an early exponential growth phase, but at stationary/senescence phase under nutrient-limiting conditions, they are assumed to be increasingly dominated by larger molecules (i.e., high molecular weight (HMW) compounds) (Borchard and Engel, 2015), and directed to DOM_L_. Mortality of all organisms, i.e., phytoplankton, bacteria and proto-zooplankton is represented by lysis (L) and death of cells (D). The latter represents cells losing their function, but preserving their integrity (i.e., apoptosis Bidle, 2016), whereby the dead biomass is directed to the small detritus pool, Det_S_, while the lysed biomass is directed towards DOM_L_, as the compounds within the cell are mainly large molecules. For phytoplankton, lysis rate is assumed to be dependent on the nutrient stress, given that it can cause reduced ability of diatoms to prevent dissolution of their cell walls (Bidle and Azam, 1999; Martin-Jézéquel et al., 2000) and increased viral mortality (Kranzler et al., 2019).

Free-living bacteria (Bac_F_) are assumed to ingest (I) exclusively DOM, whereas the particle-attached bacteria (Bac_A_) feed on detritus and TEP, the latter being a less preferred substrate (Passow, 2002). Heterotrophic flagellates (HF), in turn ingest these two bacterial groups (Massana et al., 2009), and as slightly less preferred prey, colloidal matter (Moustaka-Gouni et al., 2016), corresponding to Det_S_ and TEP in our model. Finally, ciliates feed mainly on HF, but also on bacterial groups and Det_S_ as less preferred items. In order to account for the physiological plasticities in feeding behavior (Kiørboe, 2011), and/or potential changes in community composition in response to the changes in availability of different substrate or prey items (Massana et al., 2009), initially prescribed preferences of all heterotrophs are continuously re-adjusted based on the relative concentration of different prey items, thus, reducing the importance of specified initial preferences. All heterotrophs excrete the unassimilated (U) portion as DOM and DIM. The excreted DOM by heterotrophs is assumed to be in smaller size class Buchan et al. (2014); Benner and Amon (2015), such that the function of heterotrophs, foremost the bacteria in the model is to chop the organic material into smaller pieces.

Motivated by the observation that macro-aggregates were formed temporarily after the senescence of the diatom bloom (Fig. SI-3), we explicitly described various aggregation processes in the model.The DOC molecules can coalesce (C, Passow et al., 1994) following a simplified size-based model (as in Engel et al., 2004) and form transparent exopolymer particles (TEP), which are considered to be large enough that they are retained in filters, such that they contribute to the Particulate Organic Material (POM) pool (Azam and Malfatti, 2007). TEP’s degrade (R) back to DOC_L_ at a constant specific rate (as in Schartau et al., 2007). TEP increase the stickiness, hence flocculation (F) rate of phytoplankton and small detritus (Passow, 2002), here represented as formation of large detritus, Det_L_. These flocs were assumed to leak out of the system (K), motivated by the observation that some of the debris accumulated at the bottom of the tanks was trapped inside an approximately 12cm gap at the outflow of the mesocosms, which could not be resuspended again.

We identified 8 model parameters, which were difficult to adopt from previous literature or the available observations, and to which the model results seemed sensitive. We used an objective optimization algorithm to fit these parameters. The details of the fitting procedure can be found in Section SI-1.3.

### 2.3 Numerical scenario analysis

In a first set of scenarios, we inspect the isolated impacts of a set of mechanisms on the TEP formation and POM accumulation. These are:

- ‘ExudS’: Diatom exudates are only in small DOC form, and a switch towards large DOC does not occur (Section SI-1.1.2).
- ‘noNSiL’: nutrient-stress induced increase in diatom lysis rates is turned off, so that lysis rates are maintained at low, background levels (Section SI-1.1.2).
- ‘CoagL’: Only large, and no small DOC molecules are allowed to coagulate (Section SI-1.1.4).

In a final, ‘Comb’ scenario, we analyze the effects of these three processes in combination, by combining the scenarios noNSiM, ExudS and CoagL.

## 3 Results

### 3.1 Model skill

During the development and senescence of the phytoplankton bloom (i.e., until ca. 10^*th*^ day of the experiment, corresponding to 31^*st*^ March, see chl-*a* in Fig. SI-2), consumption of nutrients is realistically reproduced by the reference model, indicating realistic description of phytoplankton growth and nutrient utilization (Fig. 2). During the last few days of the experiment, DIP and 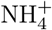 (Fig. SI-2) concentrations are overestimated, which implies either too rapid remineralization (and/or too slow nitrification for the case of 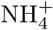), or insufficient consumption. The latter is a likely reason, given that in reality, the ability of inorganic nutrient consumption of bacteria (Pengerud et al., 1987), is not represented in our model.

**Figure 2.**
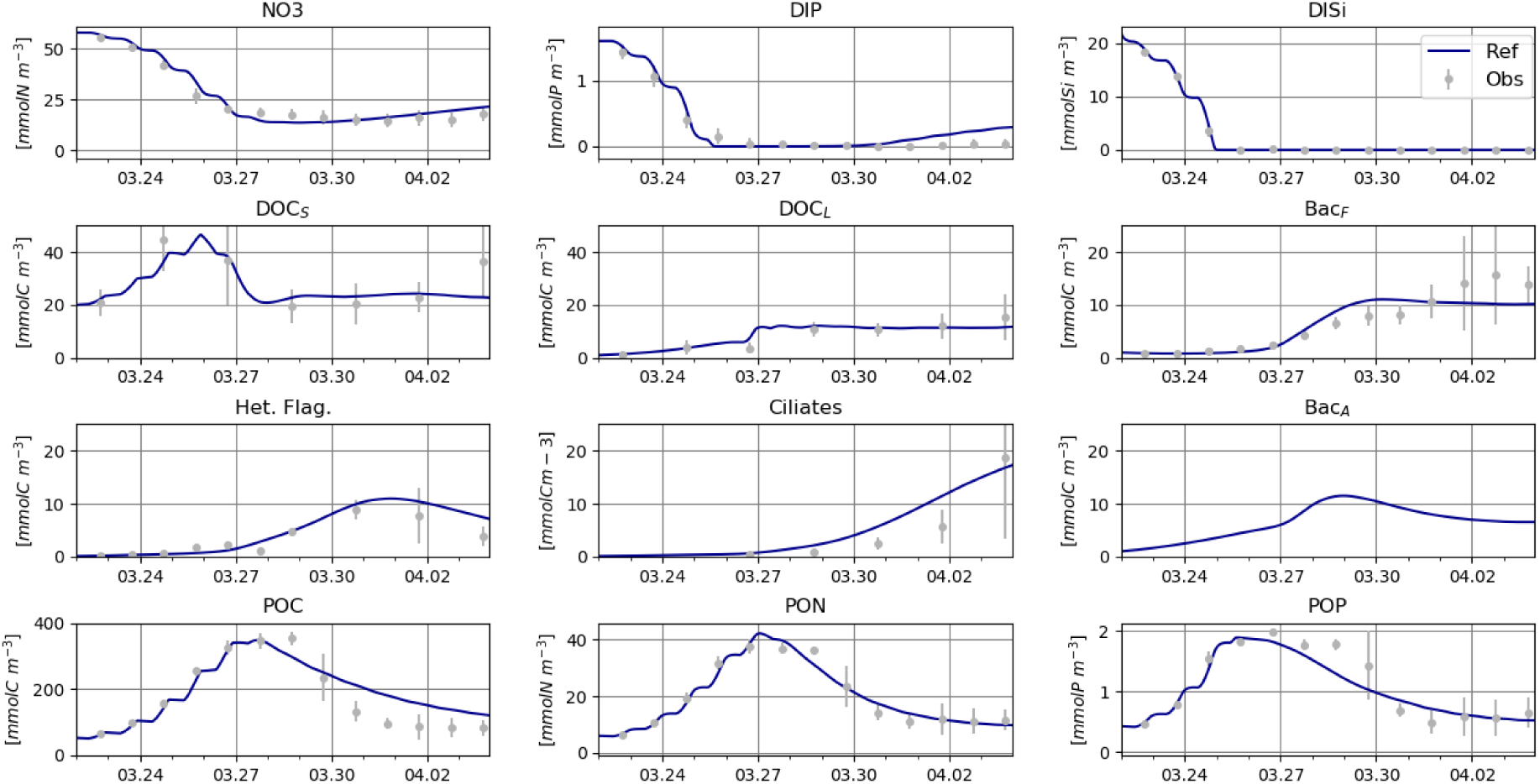
Observed (gray dots and error bars that respectively represent the mean and standard deviation across the replicates) and simulated (lines) variables considered in model calibration, except for Bac_*A*_, for which no observation data is available.

Both the DOC_S_ and DOC_L_ concentrations are in general well reproduced, except at the very end of the experiment, where the model somewhat underestimates the DOC_S_ concentrations (Fig. 2). Simulated C biomass of Bac_F_, heterotrophic flagellates and ciliates match well with the estimates based on counts, albeit some slight overshooting of protists and undershooting of Bac_F_ within the last few days of the experiment. No measurements were available to compare against the simulated Bac_A_ biomass, which was of comparable magnitude to that of *Bac*_F_ biomass, but with a slightly earlier peak, and lower concentrations towards the end. This general pattern seems to be consistent with other studies showing a switch from FL to PA bacteria during diatom blooms (Grossart et al., 2006). The increase in POC, PON and POP during the growth of diatoms, their peak concentrations, timing of their collapse, and to a lesser degree, the extent of their destruction are realistically reproduced by the model (Fig. 2).

All in all, the model can explain 90.5% of the total variation exhibited by the observations shown in Fig. 2 (see Table SI-6 for a detailed breakdown), which, to a large extent capture the cycling of C, and to a lesser extent, N and P.

### 3.2 Phytoplankton loss processes, their inter-dependencies, and consequences on material cycling

Diatom losses are dominated greatly by aggregation, followed by (total) exudation, lysis and cell death, whereas losses due to respiration constitute a minor fraction of the losses (Fig. SI-5). Exudated DOC molecules are in small form during the exponential growth phase, but become larger (Fig. SI-5) as the internal P and Si quotas start to decrease (not shown). Lysis rate of diatoms is negligible during the initial stage, but increases to considerable levels during the senescence phase (Fig. SI-5) as the cells become severely Si-limited.

Aggregation losses are caused by a steep accumulation of TEP towards the senescence phase of diatom growth, as the DOC_L_ released through exudation and lysis induced by nutrient stress rapidly form TEP, which in turn can react with the DOC_S_ that were released during the exponential growth phase, and form TEP (Fig. SI-6). It is worth noting that the contribution of the DOC_L_-DOC_L_ interaction to the TEP formation is minor, in comparison to that of the DOC_S_-TEP and the DOC_L_-TEP interactions, which are of similar magnitude (Fig SI-6). Concentration of flocs (as represented by Det_L_) increases following the diatom bloom, which is consistent with the qualitative macro-observations (Fig. SI-3), but decrease again as they are consumed by the bacteria (Bac_A_ and leak out from the system (Fig. 2).

In order to assess the relevance of some particular processes for the TEP formation, which consequently leads to a high amount of aggregation losses as explained above, we performed a numerical scenario analysis, where we analyzed the isolated and combined effects of these processes on the behavior of the system. Among the processes considered, the switch from small to large molecules in diatom exudates (‘ExudS’ scenario), and the nutrient-stress induced lysis (‘NsiL’ scenario), when considered in isolation, turned out to have negligible impact on the overall dynamics (Fig. 3). In contrast, coagulation of small DOC molecules with TEPs (‘CoagL’ scenario) seemed to have a considerable impact on a number of key variables, and the overall behavior of the system. Without this process, both the DOC_S_ and DOC_L_ concentrations reached to very high concentrations, and TEP formation, and consequently the aggregation losses and the crash of the diatom bloom were significantly delayed (Fig. 3).

**Figure 3.**
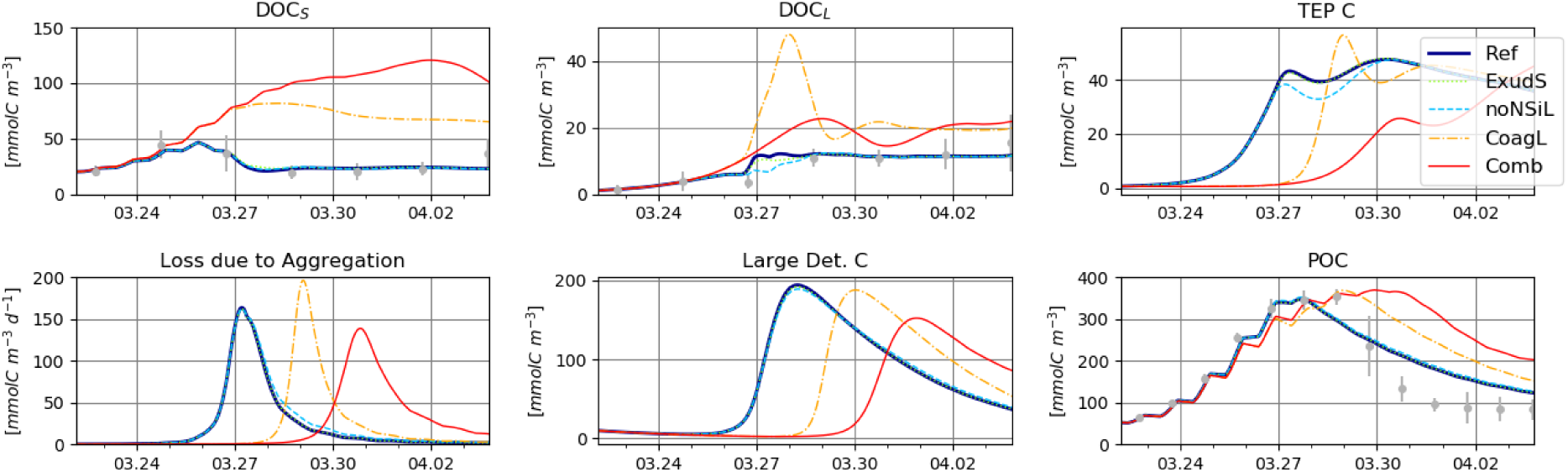
Comparison of DOC_S_, DOC_L_, TEP^C^, aggregation loss rate of phytoplankton, 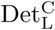 and POC, as estimated by the reference model and the three scenarios: ‘noNSIM’: lysis rate is not enhanced by nutrient stress; ‘ExudS’: exudation is only in DOC_S_ form; ‘CoagL’: only DOC_L_ (and not DOC_S_) is assumed to coagulate; ‘Comb’: where all three scenarios are combined.

Given the insignificant effects suggested by the ‘ExudS’ and ‘noNSiL’ scenarios, it could be expected that their combination with the ‘CoagL’ scenario would lead to an almost identical system behavior as in the CoagL scenario. However, this is not the case. Under the ‘Comb’ scenario, TEP formation and flocculation processes are significantly delayed, and reduced in comparison to the ‘CoagL’ scenario (Fig. 3). This result suggests synergistic interactions between these processes.

Removing the nutrient-stress dependence of lysis rates and assuming that the exudation is only in small DOC form, as represented by the ExudS scenario, DOC_L_ production by phytoplankton slows down considerably, such that their concentrations become less than in the CoagL scenario (Fig. 3). However, as DOC_S_ cannot react with TEP to produce more TEP, coagulation losses of DOC_L_ decreases even more, such that the DOC_L_ concentrations effectively becomes higher than in the Ref scenario (Fig. 3). Meanwhile DOC_S_ concentrations increase more than in the CoagL scenario, as the diatoms exudates keep being channeled to this pool. As the TEP formation is delayed more than that in the CoagL scenario, so does the flocculation processes and the crash of the diatom bloom (Fig. 3).

When CoagL is combined with ExudS, a slight deviation from the CoagL scenario occurs Fig. SI-7. Combining the CoagL with noNSiL, leads to a much more considerable difference from the CoagL scenario, suggesting that the nutrient-stress induced lysis is relatively more important than the switch from small to large DOC exudation for the behavior of the model Fig. SI-7. But the difference between the noNSiL+C and Comb scenarios is larger than the difference between the ExudS+C and Ref scenarios Fig. SI-7, pointing again to non-additive, synergistic effects.

Cumulative bulk C influxes (i.e., sum of influxes from all source terms; see Fig. SI-8 for individual fluxes driven by specific processes) to each component of the system, as simulated by the reference model and the Comb scenario exhibit significant differences especially after the first few days of the experiment (Fig. 4a). At the beginning of the experiment, C uptake by phytoplankton is the only major flux term in both model realizations. In the reference model, the build up of first DOM_S_ fueled by phytoplankton growth, and then DOM_L_ leads to an increasingly faster formation of TEP, which in turn leads to a rapid collapse of phytoplankton, and formation of flocs, i.e., Det_L_. Subsequently, the generated organic material is processed by growing bacterial subcommunites (Bac_A_, Bac_F_), and later protozooplankton. In the Comb scenario, on the other hand, exudation only in the form of DOM_S_, no enhancement in lysis rates, and DOM_S_ not interacting with TEP to form more TEP leads to trapping of most of the organic material in the form of DOM_S_. This in turn leads to significant delays in TEP formation, and consequently growth of other heterotrophs.

**Figure 4.**
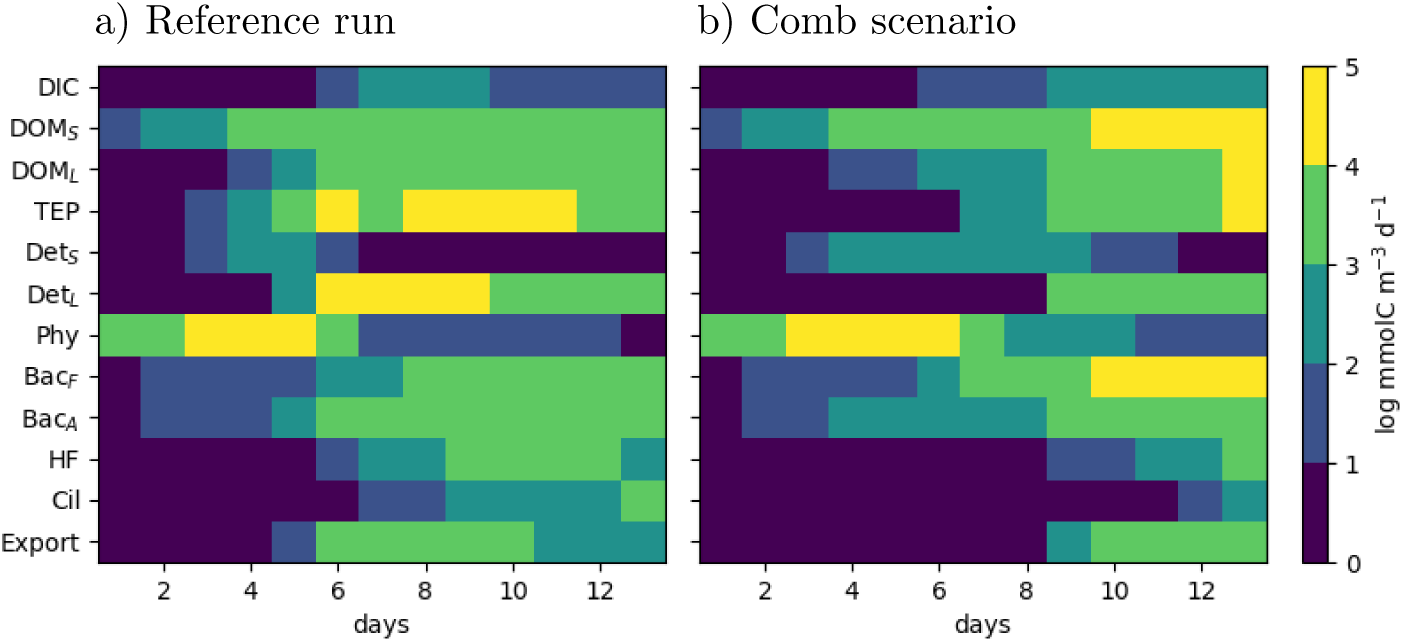
Temporal course of cumulative influxes to each model component, as simulated by a) the reference run; b) the Comb scenario, where exudation is only in DOC_S_ form, lysis rate is not enhanced by nutrient stress, and DOC_S_ does not attach to TEP.

## 4 Discussion

### 4.1 Model construction and skill

Various loss processes contributing to the termination of algal blooms may have different consequences on the organic material turnover rates (Bidle, 2016), and hence, on the post-bloom dynamics, not only within the specific context of mesocosm experiments under investigation, but also in the real marine and inland aquatic ecosystems. Therefore it is important to develop a consistent understanding of the relevance of these processes, including the specific mechanisms involved and potential dependencies and interactions between them. To this end, we developed a mechanistic numerical model to gain insight into the mechanisms governing a diatom bloom obtained in a mesocosm experiment.

Some of the processes included in the model are admittedly not readily constrainable with the available observations for this particular experiment, however they are based on qualitative evidence from earlier experimental and theoretical work. Splitting the DOC in two size classes was based on the notion that HMW and LMW compounds differ in terms of both their production and consumption by bacteria (Amon and Benner, 1996; Arnosti et al., 2018; Polimene et al., 2018). For instance, phytoplankton have been observed to release smaller molecules during the growth stage, and larger molecules during the stationary phase (Buchan et al., 2014), which we phenomenologically described in our model, in the veins of (Flynn et al., 2008; Livanou et al., 2018). Coagulation of DOC_L_ with other DOC_L_ and TEPs is similar to Engel et al. (2004), who proposed that TEP production constitutes a sink term for polysaccharides. For our system, we found that DOC_S_ also needs to coagualate with TEP, as otherwise their simulated concentration would considerably overshoot the measurements. As an alternative, we attempted to reduce their production rate (eg., by setting a different threshold composite quota value for an earlier switch to exudating larger molecules), but in that case the model was undershooting the DOC_S_ concentrations. In reality, the attachment probability of a particle depends on its own size, and the size of all other molecules in the environment, such that a size-resolved model is required to capture the interactions (Jackson, 2001). Ruiz et al. (2002) showed that a two-size class model can capture the main characteristics of a fully size resolved model, which was later employed by Engel et al. (2004) and Schartau et al. (2007). In our case we extend this to a 3-size class model, but without comparing to a full model with multiple size classes.

Initial depletion of nutrients, rise and decline of POC, PON and POP, the course of DOC concentrations, and estimated biomass of free-living bacteria and protists are well captured by the model. However DON, 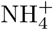 and to a lesser extent, DIP concentrations at a later phase of the experiment, are overestimated, which is presumably related with the coarse resolution of stoichiometric regulation of heterotrophs. Lack of certain data, such as the counts of attached bacteria, presumed but not quantified export (leak) of detrital material out of the system, potential diatom viruses, and size spectra of the particulate material posed challenges in validating the specific processes. Bridging these observational gaps by dynamical estimations for the corresponding variables, the model can be considered to provide a plausible representation of the mesocosm experiments during the diatom bloom and its collapse, and as such, it can be used to gain insight into the processes involved in the cycling of C especially.

### 4.2 Mechanisms contributing to the crash of diatoms

In the field, often the zooplankton is assumed to contribute greatly to the termination of phytoplankton spring blooms (e.g., Wiltshire et al., 2008; Cloern, 2018). In the planktotron experiments we presented in this study, only heteroflagellates and ciliates were present, which are too small for feeding on the large diatom species, foremost the *Thalassiosira sp*. and *Thalassiosira decipiens* that majorly contributed to the diatom bloom. Therefore zooplankton grazing can be ruled out as a potential mechanism for the crash of this particular diatom bloom.

Increasing lytic losses with nutrient stress can be caused by an increased vulnerability to bacterial (Bidle and Azam, 1999) or viral attacks (Kranzler et al., 2019), due to the compromised ability of phytoplankton to coat their cell walls with protective substances. The effect of nutrient stress on the lytic loss rates is phenomenologically accounted for in the model, without describing the exact mechanism. There exists a large amount of uncertainty regarding the loss rates caused by these processes. It is difficult to constrain the maximum rates of lysis and exudation, as it is typically not straightforward to distinguish between the DOM released through these processes (but see Ma et al., 2018; Kuhlisch et al., 2021).

Our sensitivity analysis (Section 3.2) suggests that aggregation losses are not much altered by individual processes like the changes in the fraction of small DOM in phytoplankton exudates, or the nutrient-stress induced lysis rates. However, when combined together, these processes dramatically change the TEP formation, and in turn, aggregation loss rates. This is qualitatively in agreement with field observations (e.g., Vardi et al., 2012; Lehahn et al., 2014) and experimental findings (e.g., Vincent et al., 2021).

This points to the need for further research and experimental data to be able to more precisely constrain numerical models. The specific loss processes contributing to the termination of phytoplankton blooms have often been studied in isolation so far, and our results highlights the need to consider the links between these processes, as was previously pointed out by, e.g., Vardi et al. (2012); Laber et al. (2018). From a model complexity evaluation standpoint, our results point to the challenge that evaluation of the relevance of a process description based on the typical practice of incremental alteration of model structures (e.g., Ward et al., 2013) may miss potentially relevant processes, since the combined effects of two processes can become more significant than their effects in isolation.

### 4.3 Implications and Outlook

The exact combination of the loss processes may have different implications for the post-bloom dynamics. The behavior of the system after the bloom may be sensitive to small differences in the state at bloom termination. For instance, the mesocosm duplicates (cf Section 2.1) considerably diverge from each other, beginning from around day 15: in some planktotrons, a strong, *Phaeocystis* dominated bloom developed, whereas in others, a weaker bloom was observed (Mori et al., 2021, Dlugosh et al., in prep.). Our results suggest that differences in diatom loss processes can lead to qualitative alteration in system behavior during the post-bloom phase (Figs. 3-4). Analysis of the link between the two phenomena can be assisted by our model, but after certain amendments, such as a more realistic description of algae-bacteria interactions through the production and processing of aggregates (Grossart et al., 2006).

Differences between the consequences of these processes may be upscaled in natural pelagic environments: for instance, a bloom termination predominantly governed by aggregation losses may lead to a rapid settling of OM out of the euphotic zone, and therefore clearing of the water column before the emergence of efficient zooplankton consumers, copepods in marine systems and *Daphnia* in freshwater systems. On the other hand, dominance of lytic losses as induced by viral infections can result in recycling of OM within the surface layers, and hence, maintenance or regrowth of phytoplankton biomass driven by other species, which may not be hosted by the existing viruses that had infected the species that were dominant during a previous bloom (i.e., ‘kill the winner’ Winter et al., 2010). In a mesocosm, differences in the relative importance of these mechanisms may be manifested in DOM/POM ratios; but due to the absence of vertical heterogeneities and export out of the system to a fuller extent, these post-bloom consequences are arguably not as strongly differing as can be expected in the field. Having been implemented in FABM (Bruggeman and Bolding, 2014), the model can be easily coupled with a hydrodynamical model to address these questions.

## 5 Conclusions

We developed a numerical, process-based model to analyze a spring bloom obtained in a mesocosm experiment. The model can capture *>* 90% of the total variation contained in 11 independent variables over the development and termination of the bloom. According to the simulated fluxes, aggregation losses play the most important role in the collapse of the diatom bloom, followed by exudation and nutrient-stress induced lytic losses. Coagulation of small DOC molecules with TEPs (as oppposed to coagulation of only large DOC molecules) makes a considerable difference both in DOC concentrations, and TEP formation, hence the timing of the bloom termination. Switches in the form of exudated DOC (i.e., small to large), and the lytic losses have negligible effects when considered in isolation, however, when combined with coagulation possibility of small DOC molecules, they make a significant difference, hence, pointing to synergistic interactions. These results have ramifications on the development of complex biogeochemical models, and the present understanding of the coupling between aggregation, microbial and plankton networks.

## Acknowledgments

We are grateful to all the other EcoMol Planktotron team members, not listed here as co-authors but whose efforts made the experiment possible and successful: Vera Bischoff, Simon Dufner, Bert Engelen, Lena Engelmann, Ferdinand Esser, Oliver Ferdinand, Ulrike Feudel, Jan Freund, Matthias Friebe, Andrea Gall, Hans-Jürgen Brumsack, Bernhard Schnetger, Katharina Pahnke, Helge-Ansgar Giebel, Eleonore Grundken, Charlotta Hecker, Birgit Kürzel, Carola Lehners, Raquel Lopes, Julian Merder, Felix Milke, Cristina Moraru, Lars-Eric Peterson, Christoph Plum, Ralf Rabus, Heike Rickels, Vanessa Schnaars, Matthias Schroeder, Peter Schupp, Ina Ulber, Marie von Schneden, Rolf Weinert, Lars Woehlbrand, Mathias Wolterink and Oliver Zielinski. We would like to further thank the captain and crew of the RV Heincke, and the ICBM machine shop. We thank Tropic Marine (Hünenberg, Switzerland), who kindly provided us with the Tropic Marine Pro-Reef Sea Salt required for the artificial seawater medium. Finally we acknowledge the Lower Saxon State Department for Waterway, Coastal and Nature Conservation (NLWKN) for providing microscopic measurement data.crew of the RV Heincke, and the ICBM machine shop. We thank Tropic Marine (Hünenberg, Switzerland), who kindly provided us with the Tropic Marine Pro-Reef Sea Salt required for the artificial seawater medium. Finally we acknowledge the Lower Saxon State Department for Waterway, Coastal and Nature Conservation (NLWKN) for providing microscopic measurement data.crew of the RV Heincke, and the ICBM machine shop. We thank Tropic Marine (Hünenberg, Switzerland), who kindly provided us with the Tropic Marine Pro-Reef Sea Salt required for the artificial seawater medium. Finally we acknowledge the Lower Saxon State Department for Waterway, Coastal and Nature Conservation (NLWKN) for providing microscopic measurement data.crew of the RV Heincke, and the ICBM machine shop. We thank Tropic Marine (Hünenberg, Switzerland), who kindly provided us with the Tropic Marine Pro-Reef Sea Salt required for the artificial seawater medium. Finally we acknowledge the Lower Saxon State Department for Waterway, Coastal and Nature Conservation (NLWKN) for providing microscopic measurement data.

## Author contributions

OK and MSi conceptualized the work, OK developed the model and wrote the initial draft, OK and LL developed formal analysis tools, including validation; NHH, LB, CB, CM contributed to data curation; NHH and BH contributed to the investigation; NHH, Mst, CM, BB, MSi contributed to the writing and reviewing of the work; TD, MSt, MSi were responsible for the project administration and experimental methodology.

## Funding

OK was supported by the Deutsche Forschungsgemeinschaft (DFG, KE1970/2-1). This study was carried out in the framework of the Ph.D. research training group ‘The Ecology of Molecules’ (EcoMol) supported by the Lower Saxony Ministry for Science and Culture (MWK). The establishment of the infrastructure was financed by the MWK and the Institute for Chemistry and Biology of the Marine Environment (ICBM). This work was partially supported by the Collaborative Research Center Roseobacter (TRR51) funded by Deutsche Forschungsgemeinschaft. Funding to CB was generously provided by the Helmholtz Institute for Functional Marine Biodiversity (HIFMB), a collaboration between the Alfred-Wegener-Institute, Helmholtz-Center for Polar and Marine Research, and the Carl-von-Ossietzky University Oldenburg, initially funded by the MWK and the Volkswagen Foundation through the ‘Niedersächsisches Vorab’ grant program (grant number ZN3285). The inoculum for the mesocosms was derived during RV Heinke research cruise HE504 (grant number AWI HE504 leg2 00).

## Notes

### Competing Interest Statement

The authors have declared no competing interest.

