## Supporting Information for "Growth, organic matter release, aggregation and recycling during a diatom bloom: A model-based analysis of a mesocosm experiment"

### SI-1 Details of the Model

The model used in this study is based on the pelagic modules of the 'Generalized Plankton Model' (GPM, Kerimoglu et al., 2020). GPM was adapted to the purposes of the current study, by:

- considering a single phytoplankton type, i.e., the diatoms, given that the biomass of all other groups remain low until the termination of the diatom bloom.
- refining the representation of silicate uptake and co-limitation
- describing two DOM pools for small (i.e., $DOM_S$, monomers and low molecular weight compounds) and large ($DOM_L$, high molecular weight compounds) instead of a single DOM pool.
- refining the targets of loss terms (exudation, lysis, death) of phytoplankton.
- describing the formation of transparent exopolymer particles (TEPs), through coalescence of DOM.
- describing the aggregation of diatoms, small detritus particles and TEPs into flocs, represented here by large detritus
- describing the dynamics of free living and particle-attached bacteria ($Bac_F$ and $Bac_A$, respectively), their ingestion (of DOM, detritus and TEPs), and excretion of unassimilated portion in the form of DOM and DIM

- describing the dynamics of heterotrophic flagellates (HF), and ciliates (Cil)
- refining the representation of silicate remineralization.

A full description of processes included in the model (Section SI-1.1), its implementation for simulating the mesocosms (Section SI-1.2), and calibration (Section SI-1.3) are provided below.

#### SI-1.1 Process Descriptions

In Section SI-1.1.1, we first provide an overview of the system of Ordinary Differential Equations (ODEs), and the source-sink terms governing the dynamics of material present in each pool (Fig. 1). Then in the following sections (SI-1.1.2 – SI-1.1.5), detailed description of individual processes are given for each model component.

##### SI-1.1.1 ODE System

Phytoplankton biomass is represented by 5 state variables for C, Chl, P, N and Si bound to the organisms, in order to account for the variability in Chl:C:N:P:Si ratio of phytoplankton. Source and sink terms of each of these state variables are as follows:

$$\dot{Phy}^{\mathrm{C}} = \underbrace{P^{\mathrm{C}}}_{\text{Production}} - \underbrace{R_P^{\mathrm{C}}}_{\text{Respiration}} - \underbrace{E^{\mathrm{C}}}_{\text{Exudation}} - \underbrace{L_P^{\mathrm{C}}}_{\text{Lysis}} - \underbrace{D_P^{\mathrm{C}}}_{\text{Death}} - \underbrace{A2_P^{\mathrm{C}}}_{\text{Aggr.-2}} \tag{SI-1a}$$

$$\dot{Phy}^{\mathrm{Chl}} = \underbrace{V^{\mathrm{N}}\rho^{\mathrm{Chl}}}_{\text{Synthesis}} - \underbrace{R^{\mathrm{Chl}}}_{\text{Degradation}} - (L_P^{\mathrm{C}} + D_P^{\mathrm{C}} + A2_P^{\mathrm{C}})\theta \tag{SI-1b}$$

$$\dot{Phy}^{\mathrm{X}} = \underbrace{V^{\mathrm{X}}}_{\text{X uptake}} - E^{\mathrm{X}} - (L_P^{\mathrm{C}} + D_P^{\mathrm{C}} + A2_P^{\mathrm{C}})Q_P^{\mathrm{X}},\ X = \{N, P, Si\} \tag{SI-1c}$$

where, $Q_P^{\mathrm{X}}$ are the quotas of X={N, P, Si} expressed as their molar ratio to the C content, and $\theta$ is the Chl:C ratio of phytoplankton. Individual terms are explained in setion SI-1.1.2.

For being able to better track the TEP formation, which mostly consist of the large C-rich polysaccharides (Passow, 2002) we split the DOC pool into two, that represent small ($DOC_S$) and large

$(DOC_L)$ molecules:

$$\dot{DOC_S} = E^{\mathrm{C}} f_S^E + \underbrace{\sum_i \sum_j I_{i,j}^{\mathrm{C}} (1-\epsilon_i^{\mathrm{C}})\, f_i^{UO}}_{\text{Unas. Ing.}} - \underbrace{\sum_i I_{i,DOC_S}}_{\text{Ing. loss}} - \underbrace{A1_{DOC_S}^{\mathrm{C}}}_{\text{Aggr.-1}} \tag{SI-2a}$$

$$\dot{DOC_L} = E^{\mathrm{C}} (1-f_S^E) + L_P^{\mathrm{C}} + \sum_i L_i^C + \underbrace{R_{TEP}}_{\text{Degr.}} - \sum_i I_{i,DOC_L} - \underbrace{A1_{DOC_L}^{\mathrm{C}}}_{\text{Aggr.-1}} \tag{SI-2b}$$

$$\dot{DOX} = E^{\mathrm{X}} + L_P^{\mathrm{C}} Q_P^{\mathrm{X}} + \sum_i L_i^C Q_i^{\mathrm{X}} + \sum_i \sum_j I_{i,j}^{\mathrm{X}} (1-\epsilon_i^{\mathrm{X}}) - \sum_i I_{i,DOX}, \; X = \{N, P\} \tag{SI-2c}$$

where $f_S^E$ describes the partitioning of DOC exudates of phytoplankton to large and small classes (Section SI-1.1.2); $\epsilon^{\mathrm{C}}$ and $f_i^{UO}$ respectively describe the assimilation efficiency and partitioning of unassimilated N and P excretions of heterotrophs to DOM and DIM pools (Section SI-1.1.3); and finally $i$ and $j$ index the heterotrophs, and their prey targets, respectively (Section SI-1.1.3); A similar splitting of DOC pool into the small and large fractions was employed by Flynn et al. (2008), for a more accurate description of phytoplankton - DOM fluxes. Carbon bound to Transparent Exopolymeric Particles (TEP) is explicitly tracked with a state variable:

$$\dot{TEP}^{\mathrm{C}} = \underbrace{A1_{DOC_S}^{\mathrm{C}} + A1_{DOC_L}^{\mathrm{C}}}_{\text{Aggr.-1}} - R_{TEP} - A2_{TEP}^{\mathrm{C}} - \sum_i I_{i,TEP}^{\mathrm{C}} \tag{SI-3}$$

Two detritus size classes are considered: the 'small' detritus size class represents the freshly dead phytoplankton and bacteria.

$$\dot{Det}_S^{\mathrm{C}} = D_P^{\mathrm{C}} + \sum_i D_i^{\mathrm{C}} - \sum_i I_{i,Det_S}^{\mathrm{C}} - A2_{Det_S}^{\mathrm{C}} \tag{SI-4a}$$

$$\dot{Det}_S^{\mathrm{X}} = D_P^{\mathrm{C}} Q_P^{\mathrm{X}} + \sum_i D_i^{\mathrm{C}} Q_i^{\mathrm{X}} - \sum_i I_{i,Det_S}^{\mathrm{X}} - A2_{Det_S}^{\mathrm{X}}, \; X = \{N, P\} \tag{SI-4b}$$

$$\dot{Det}_S^{\mathrm{Si}} = D_P^{\mathrm{C}} Q_P^{\mathrm{Si}} + \sum_i D_i^{\mathrm{C}} Q_i^{\mathrm{Si}} + \sum_i \sum_j I_{i,j}^{\mathrm{Si}} f_i^{UsiD} - \sum_i I_{i,Det_S}^{\mathrm{Si}} - A2_{Det_S}^{\mathrm{Si}} \tag{SI-4c}$$

where, the last term in $Det_S^{\mathrm{Si}}$ represents the non-dissolved fraction of the uningested Si with $f_i^{UsiD}$ describing the detrital fraction, rest being DIM (Eq. SI-7e).

The 'large' detritus size class represents the flocs that mainly form by the aggregation of small detritus, phytoplankton and TEPs:

$$\dot{Det}_L^{\mathrm{C}} = A2_P^{\mathrm{C}} + A2_{Det_S}^{\mathrm{C}} + A2_{TEP}^{\mathrm{C}} - \sum_i I_{i,Det_L}^{\mathrm{C}} - \underbrace{Det_L^{\mathrm{C}} k_l}_{\text{Leakage}} \tag{SI-5a}$$

$$\dot{Det}_L^{\mathrm{X}} = A2_P^{\mathrm{X}} + A2_{Det_S}^{\mathrm{X}} - \sum_i I_{i,Det_L}^{\mathrm{X}} - Det_L^{\mathrm{X}} k_l, \;\; X = \{N, P, Si\} \tag{SI-5b}$$

where, the leakage term describes the export of flocs out of the system, as motivated by the macroscopic observation that flocs were falling inside the instrumentation gap at the bottom of the tanks. We assumed a conservative rate of $k_l = 0.1$ $\mathrm{d}^{-1}$ (Table SI-4).

Biomass of heterotrophs considered in this study, i.e., the two bacterial groups, $\mathrm{Bac_A}$ and $\mathrm{Bac_F}$, and two planktonic protists, HF and Cil are represented by their C content only, the variability in the C:N:P ratio are ignored:

$$\dot{Het}_i^{\mathrm{C}} = \underbrace{\sum_j I_{i,j}^{\mathrm{C}} \, \epsilon_i^C}_{\text{Ing. gain}} - \underbrace{\sum_i I_{i,j}^{\mathrm{C}}}_{\text{Ing. loss}} - \underbrace{R_i^{\mathrm{C}}}_{\text{Respiration}} - \underbrace{L_i^{\mathrm{C}}}_{\text{Lysis}} - \underbrace{D_i^{\mathrm{C}}}_{\text{Death}} \;\; i = \{Bac_A, Bac_F, HF, Cil\} \tag{SI-6}$$

Dissolved Inorganic Material (DIM) consists of DIC (only for tracking the mass balance), two forms of inorganic nitrogen, nitrate ($\mathrm{NO_3^-}$) and ammonium ($\mathrm{NH_4^+}$), phosphorus (DIP) and silicate (DISi):

$$\dot{DIC} = -P^{\mathrm{C}} + R_P^{\mathrm{C}} + R_H^{\mathrm{C}} + \sum_i \sum_j I_{i,j}^{\mathrm{C}} (1 - \epsilon_i^{\mathrm{C}})(1 - f_i^{UO}) \tag{SI-7a}$$

$$\dot{DIP} = -V^{\mathrm{P}} + \sum_i \sum_j I_{i,j}^{\mathrm{P}} (1 - \epsilon_i^{\mathrm{P}})(1 - f_i^{UO}) \tag{SI-7b}$$

$$\dot{NH_4^+} = -V^{\mathrm{NH4}} - \underbrace{Z}_{\text{Nitrification}} + \sum_i \sum_j I_{i,j}^{\mathrm{N}} (1 - \epsilon_i^{\mathrm{N}})(1 - f_i^{UO}) \tag{SI-7c}$$

$$\dot{NO_3^-} = -V^{\mathrm{NO3}} + Z \tag{SI-7d}$$

$$\dot{DISi} = -V^{\mathrm{Si}} + E^{\mathrm{Si}} + \sum_i \sum_j I_{i,j}^{\mathrm{Si}} (1 - f_i^{UsiD}) \tag{SI-7e}$$

##### SI-1.1.2 Phytoplankton

Bulk production rate of phytoplankton is calculated as:

$$P^{\mathrm{C}} = Phy^{\mathrm{C}} \underbrace{\underbrace{\mu_{max}\mathscr{F}_{\mathrm{T}}\mathscr{F}_{\mathrm{nut}}}_{p_m^{\mathrm{C}}}\mathscr{F}_{\mathrm{par}}}_{p^{\mathrm{C}}} \tag{SI-8}$$

where, $\mathscr{F}_{\mathrm{T}}$ is the Arrhenius function (Eq. SI-39), $\mathscr{F}_{\mathrm{nut}}$ is the overall nutrient limitation function, $\mathscr{F}_{\mathrm{par}}$ is the irradiance limitation function and the $p_m^{\mathrm{C}}$ is the light-saturated specific production rate and $p^{\mathrm{C}}$ is the specific production rate. $\mathscr{F}_{\mathrm{nut}}$ describes the co-limitation by N, P and Si according to the Liebig's law of minimum:

$$\mathscr{F}_{\mathrm{nut}} = \min\left(\mathscr{F}_P^{\mathrm{P}}, \mathscr{F}_P^{\mathrm{N}}, \mathscr{F}_P^{\mathrm{Si}}\right) \tag{SI-9}$$

The individual limitation factors for $\mathscr{F}_P^{\mathrm{X}}$ (X={N,P,Si}) are calculated as functions of their internal availability within the cell based on the Caperon/Droop approach (Caperon, 1968; Droop, 1968):

$$\mathscr{F}_P^{\mathrm{X}} = \left(1 - \frac{Q_{P,min}^{\mathrm{X}}}{Q_P^{\mathrm{X}}}\right),\ \mathrm{X} = \{\mathrm{N,P,Si}\} \tag{SI-10}$$

Based on the observation that Si uptake and division rates for diatoms are highly coupled (Martin-Jézéquel et al., 2000), it was suggested that Si limitation can be better described by the transport rate, regardless of the cellular Si:C ratio (e.g., Flynn and Martin-Jézéquel, 2000). We tested this approach, but it lead to plateauing of diatom biomass and chlorophyll concentrations as soon as the external Si got depleted, which is not supported by the observations obtained in the experiments under consideration (see Fig.x and Fig.y). We found that the best formulation to reproduce the observed nutrient consumption and chlorophyll patterns was to use the standard Caperon/Droop approach for Si, but use a relatively high subsistence quota (see Table SI-1) such that growth is not immediately ceased, but soon after the depletion of external Si.

$\mathscr{F}_{\mathrm{par}}$, describes the saturation of photosynthesis with increasing photosynthetically active radiation (PAR) and $\theta$, according to Geider et al. (1998):

$$\mathscr{F}_{\mathrm{par}} = 1 - \exp\left(\frac{-\alpha^{Chl}\theta\mathrm{PAR}}{P_m^{\mathrm{C}}}\right) \tag{SI-11}$$

As in Geider et al. (1998), bulk chlorophyll synthesis rate is assumed to be proportional to bulk nitrogen uptake rate, $V^{\mathrm{N}}$, and the imbalance between production and light absorption rates, $\rho^{Chl}$ (Eq. SI-1b). The latter is given by:

$$\rho^{\mathrm{Chl}} = \theta^{\mathrm{N}}_{max} \frac{p^{\mathrm{C}}}{\alpha^{Chl}\theta I} * 12\mathrm{gC/molC} \tag{SI-12}$$

where $\theta^{\mathrm{N}}_{max}$ as originally used by Geider et al. (1998) is approximated by $\theta_{max}/Q^{\mathrm{N}}_{P,max}$ in order to eliminate the parameter $\theta^{\mathrm{N}}_{max}$. Specific chlorophyll degradation rate is assumed to be the same as the phytoplankton respiration rate, i.e., $R^{\mathrm{Chl}} = R^{\mathrm{C}}_{P}\theta$.

Finally, the bulk uptake rate of nutrients, $V^{\mathrm{X}}$ (X={N,P,Si}, $V^{\mathrm{N}} = V^{NO3} + V^{NH4}$) are calculated as follows:

$$V^{\mathrm{X}} = Phy^{\mathrm{C}} \underbrace{\mathscr{F}_{\mathrm{T}} v^{\mathrm{X}}_{max}(1 - \mathscr{F}_{Q^{\mathrm{X}}_{P}})\mathscr{F}_{E^{X}}}_{v^{\mathrm{X}}},\ X = \{\mathrm{P,\ Si,\ NO_3^-,\ NH_4^+}\} \tag{SI-13a}$$

where $v^{\mathrm{X}}$ and $v^{\mathrm{X}}_{max}$ (X={P,N,Si}) represent the specific, and maximum specific uptake rates, respectively, and $\mathscr{F}_{Q^{\mathrm{X}}}$ describe the relative quota of the cells:

$$\mathscr{F}_{Q^{\mathrm{X}}_{P}} = \frac{Q^{\mathrm{X}}_{P} - Q^{\mathrm{X}}_{P,min}}{Q^{\mathrm{X}}_{P,max} - Q^{\mathrm{X}}_{P,min}},\ \mathrm{X} = \{\mathrm{N,\ P,\ Si}\} \tag{SI-14}$$

with $Q^{\mathrm{X}}_{max}$ and $Q^{\mathrm{X}}_{min}$ are the prescribed maximum and minimum quotas, such that 1-$\mathscr{F}_{Q^{\mathrm{X}}}$ describes linear down-regulation of uptake rate as a function of internal quotas (Eq. SI-14). Finally, $\mathscr{F}_{E_X}$ ($X = \{\mathrm{DIP,\ NO_3^-,\ NH_4^+}\}$ describes dependence of uptake rate on the external concentration of nutrients (Eq. SI-15a):

$$\mathscr{F}_{E_X} = \frac{X}{X + K_X},\ X' = \{\mathrm{DIP,\ DISi}\} \tag{SI-15a}$$

$$\mathscr{F}_{E_X} = \frac{X'}{1 + \mathrm{NO_3^-}' + \mathrm{NH_4^+}'},\ X' = \{\mathrm{NO_3^-}',\ \mathrm{NH_4^+}'\} \tag{SI-15b}$$

where $\mathrm{NO_3^-}'$ and $\mathrm{NH_4^+}'$ are the concentrations of $\mathrm{NO_3^-}$ and $\mathrm{NH_4^+}$ weighed by their half saturation constants, i.e., $\mathrm{NO_3^-}' = \mathrm{NO_3^-}/K_{NO3}$ and $\mathrm{NH_4^+}' = \mathrm{NH_4^+}/K_{NH4}$.

Both the respiration and exudation rates are calculated as the sum of a base and a factor of growth (production) rates:

$$R_P^{\mathrm{C}} = Phy^{\mathrm{C}}\zeta_{pb} + P^{\mathrm{C}}\zeta_{pp} \tag{SI-16}$$

$$E^{\mathrm{C}} = Phy^{\mathrm{C}}\gamma_{pb} + P^{\mathrm{C}}\gamma_{pp} \tag{SI-17}$$

As long as the cellular quotas of diatoms are below the maximum specified values, the N, P and Si content of exudates, $E^{\mathrm{X}}$ (Eq. SI-1c; Eq. SI-2c) are assumed to be negligable , but as $Q_P^{\mathrm{X}}$ approaches $Q_{P,max}^{\mathrm{X}}$, $E^{\mathrm{X}}$ become proportional to $E^{\mathrm{C}}$ with $Q^{\mathrm{X}}$ being the proportionality constant. This switch is described by a piecewise Heaviside function of relative quota, $\mathscr{H}^{\mathrm{X}}$:

$$E^{\mathrm{X}} = \mathscr{H}^{\mathrm{X}} E^{\mathrm{C}} Q_P^{\mathrm{X}} \tag{SI-18}$$

where,

$$\mathscr{H}^{\mathrm{X}} = \begin{cases} 0, & \mathscr{F}_{Q_P^{\mathrm{X}}} \leq 0.99 \\ 1, & \mathscr{F}_{Q_P^{\mathrm{X}}} > 0.99 \end{cases} \quad X' = \{N, P, Si\} \tag{SI-19}$$

In reality, quality of DOM produced by phytoplankton depends on the specific phase of growth (Flynn et al., 2008). Overall, DOC-L is mainly supplied by the exudation of 'overconsumed' carbon (Toggweiler, 1993), but also by lysis of phytoplankton and bacterial cells.

Exudates are assumed to consist of smaller molecules during the exponential growth phase of diatoms, but larger molecules at the senescence phase, under nutrient stress, as suggested by (e.g., Passow, 2002; Borchard and Engel, 2015). This is implemented in the model by describing the fraction of small exudates, $f_S^E$ to sigmoidally increase from 0 to 1, i.e., from exclusively small to exclusively large DOM (see Eq. SI-2a-b) with increasing overall nutrition status, $\mathscr{F}_{nut}$ (Fig. SI-1):

$$f_S^E = (1.0 + exp(-(\mathscr{F}_{nut} - 0.2) \cdot 30))^{-1} \tag{SI-20}$$

where, $\mathscr{F}_{nut}$ is identified by the minimum of the relative N, P and Si quotas:

$$\mathscr{F}_{nut} = \min(\mathscr{F}_{Q^{\mathrm{N}}}, \mathscr{F}_{Q^{\mathrm{P}}}, \mathscr{F}_{Q^{\mathrm{Si}}}) \tag{SI-21}$$

Whole-cell loss terms, i.e., lysis, death and aggregation are calculated based on the C content of

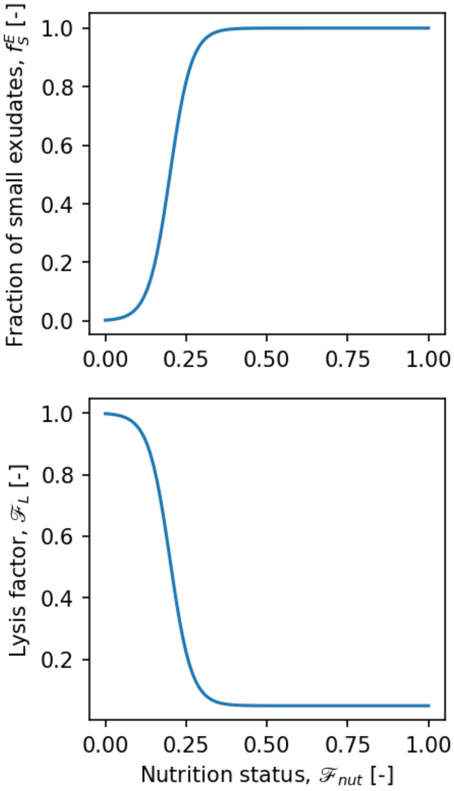


**Figure SI-1.** Fraction of small exudates and lysis factor as a function of nutrition status of cells.

phytoplankton, respective rates for N, P and Chl are then calculated based on these rates and the ratio of N, P and Chl to C of the cell (Eq. SI-1b,c,d, Eq. SI-2c,d, Eq. SI-4b,c). Aggregation, which affects non-phytoplanktonic particles as well, is described in SI-1.1.4. Diatoms have been demonstrated to be more susceptible for viral mortality under silicate limitation Kranzler et al. (2019). Moreover, under nutrient limitation, ability of phytoplankton to defend their cell walls against bacterial attacks becomes compromised (Martin-Jézéquel et al., 2000). Therefore, lysis rate is assumed to dependent on the nutrition state of the cells. This is described by sigmoidal function that sharply increases from a bacgkround fraction (assumed to be 5%) to 100% of the prescribed specific lysis rate, $l_P^l$ (Fig. SI-1):

$$L_P^{\mathrm{C}} = Phy^{\mathrm{C}}(\mathscr{F}_L l_P^l) \tag{SI-22}$$

where,

$$\mathscr{F}_L = 0.05 + 0.95\,(1.0 + exp(-(\mathscr{F}_{nut} - 0.2)\cdot 30))^{-1} \tag{SI-23}$$

Lysed phytoplankton material and exudated substances are added to the $DOM_\mathrm{L}$ pool (Eq. SI-2a). Cell death, $D_P^\mathrm{X}$ , which represents the inactivation (e.g., slowly dying) of cells (Bidle, 2016), is considered to have a linear component only.

$$D_P^\mathrm{C} = Phy^\mathrm{C} m_P^l \tag{SI-24}$$

Death losses are channeled to the small detritus class (Eq. SI-4).

**Table SI-1.** Parameters of the phytoplankton module. Sources: A: Assumed; T: Hand-Tuned C: Auto-Calibrated. G97: Geider et al. (1997) for *Thalassiosira weissflogii*; T20: Tillmann et al. (2000); G98: Geider et al. (1998); P13: Pahlow et al. (2013) for *Thalassiosira fluviatilis*; DG99: Davidson and Gurney (1999); S07: Schartau et al. (2007) referring to Ruiz et al. (2002); R12: (Regaudie-De-Gioux and Duarte, 2012); O05: Oubelkheir et al. (2005).

| Symbol | Description | Value | Unit | Source |
|---|---|---|---|---|
| $\mu_\mathrm{max}$ | Maximum growth rate | 5.2 | $d^{-1}$ | G97 |
| $\alpha^{Chl}$ | Chl. sp. slope of P-I curve | 7.0 | gC gChl$^{-1}$ / (E m$^{-2}$) | T20 |
| $\theta_{max}$ | Max. Chl:C ratio | 0.07 | m$^2$ gChl/gC$^{-1}$ | T |
| $Q^\mathrm{N}_{P,max}$ | Maximum quota for N | 0.15 | molN molC$^{-1}$ | T |
| $Q^\mathrm{P}_{P,max}$ | Maximum quota for P | 0.012 | molP molC$^{-1}$ | T |
| $Q^\mathrm{Si}_{P,max}$ | Maximum quota for Si | 0.18 | molSi molC$^{-1}$ | T |
| $Q^\mathrm{N}_{P,min}$ | Subsistence N quota | 0.05 | molN molC$^{-1}$ | G98 |
| $Q^\mathrm{P}_{P,min}$ | Subsistence P quota | 0.003 | molP molC$^{-1}$ | A |
| $Q^\mathrm{Si}_{P,min}$ | Subsistence Si quota | 0.06 | molSi molC$^{-1}$ | A |
| $v^\mathrm{N}_\mathrm{max}$ | Maximum N uptake rate | 0.78 | molN (mmolC d)$^{-1}$ | $= \mu_\mathrm{max} Q^\mathrm{N}_{P,max}$ |
| $v^P_\mathrm{max}$ | Maximum P uptake rate | 0.075 | molP (mmolC d)$^{-1}$ | T |
| $v^\mathrm{Si}_\mathrm{max}$ | Maximum Si uptake rate | 1.2 | molSi (mmolC d)$^{-1}$ | A |
| $K_{NO3}$ | Half saturation constant for $NO_3$ uptake | 2.0 | mmolN m$^{-3}$ | G98 |
| $K_{NH4}$ | Half saturation constant for $NH_4$ uptake | 0.5 | mmolN m$^{-3}$ | G98 |
| $K_{DIP}$ | Half saturation constant for P uptake | 0.05 | mmolP m$^{-3}$ | A |
| $K_{Si}$ | Half saturation constant for Si limitation | 0.43 | mmolSi m$^{-3}$ | DG99 |
| $d^l_P$ | Specific death rate | 0.05 | $d^{-1}$ | T |
| $\zeta_{pb}$ | Basal respiration rate | 0.01 | $d^{-1}$ | A |
| $\zeta_{pp}$ | Production sp. respiration rate | 0.01 | - | A |
| $\gamma_{pb}$ | Basal exudation rate | 0.02 | $d^{-1}$ | T |
| $\gamma_{pp}$ | Production sp. exudation rate | 0.11 | - | C |
| $l^l_P$ | Specific lysis rate | 0.1 | $d^{-1}$ | C |
| $E_a$ | Activation energy for temperature scaling | 0.32 | - | R12 |
| $T_{ref}$ | Reference Temperature | 283 | K | - |
| $k^\mathrm{P}_c$ | Specific attenuation coefficient | 0.024 | m$^2$ mmolC$^{-1}$ | O05 |

#### SI-1.1.3 Heterotrophs

We consider two functional groups of bacteria: the free-living bacteria, $\mathrm{Bac_F}$, which are assumed to feed on DOM (both size classes), and the particle-attached bacteria, $\mathrm{Bac_A}$, which are assumed to feed on detrital material, and TEP as a slighly less preferred item. In addition, we consider two heterotrophic protists: the heteroflagellates, HF, and ciliates, Cil. HF are assumed to feed on the two forms of bacteria as preferred items, and small detritus and TEP. Cil feed mainly on HF, but also on all items that HF can feed on as less preferred items.

For a heterotroph $H_i$, C-based bulk ingestion rate of a feeding target $t_j$ is given by:

$$I_{i,j}^{\mathrm{C}} = H_i^{\mathrm{C}} I_{i,max} \frac{pw_{i,j} t_j^{\mathrm{C}}}{K_i^I + \sum_j (pw_{i,j}) t_{i,j}^{\mathrm{C}}} \tag{SI-25}$$

Following Fasham et al. (1990), prescribed raw preferences of prey items (Table SI-2) are dynamically weighed with their relative C concentration to determine the realized preferences, $pw$:

$$pw_{i,j} = \frac{p_{i,j} t_j^{\mathrm{C}}}{\sum_j (p_{i,j} t_j^{\mathrm{C}})} \tag{SI-26}$$

Besides stabilizing model solutions, these dynamically changing preferences can be thought to simplistically represent the shifts in community composition in response to changes in substrate availability (e.g., Teeling et al., 2012, for bacteria).

**Table SI-2.** Raw preferences, $p_{i,j}$ of heterotroph $i$ for target $j$ (columns).

| $H_i \setminus t_j$ | $DOC_S$ | $DOC_L$ | $TEP^{\mathrm{C}}$ | $Det_S^{\mathrm{C}}$ | $Det_L^{\mathrm{C}}$ | $Bac_A$ | $Bac_F$ | $HF$ |
|---|---|---|---|---|---|---|---|---|
| $Bac_F$ | 0.5 | 0.5 | 0.0 | 0.0 | 0.0 | 0.0 | 0.0 | 0.0 |
| $Bac_A$ | 0.0 | 0.0 | 0.2 | 0.4 | 0.4 | 0.0 | 0.0 | 0.0 |
| $HF$ | 0.0 | 0.0 | 0.1 | 0.1 | 0.0 | 0.4 | 0.4 | 0.0 |
| $Cil$ | 0.0 | 0.0 | 0.1 | 0.1 | 0.0 | 0.1 | 0.1 | 0.6 |

Ingestion rate of N-, P-, and Si- are calculated by multiplying the C-based ingestion rate with the respective ratio of their content to the C content within the target item:

$$I_{i,j}^{\mathrm{X}} = I_{i,j}^{\mathrm{C}} \frac{t_j^{\mathrm{X}}}{t_j^{\mathrm{C}}},\ \mathrm{X} = \{\mathrm{N,P,Si}\} \tag{SI-27}$$

Assimilated portion of the ingested item $i$ is given by the product of ingestion rate and element-specific assimilation efficiency, $\epsilon^{\mathrm{X}}$ (Eq.SI-6), and unassimilated fraction is the rest (e.g., Eq.SI-2). A constant fraction of the unassimilated C, N and P, $f_i^{UO}$, is assumed to be excreted into $\mathrm{DOM_S}$ and the rest to the DIM pool (Eq. SI-7,SI-2). This fraction is assumed to be smaller for bacteria (Table SI-3, Buchan et al. (2014)). A constant, and large fraction of Si in ingested prey, $f_i^{UsiD}$, is assumed to be excreted in detritus form, and the rest to be remineralized as DIM (Eq. SI-7,SI-4). This fraction is again assumed to be slightly lower for bacteria (Table SI-3 (Bidle and Azam, 1999). The $\epsilon^{\mathrm{X}}$ are dynamically adjusted according to the difference between the stoichiometries of the ingested material and the heterotrph itself, based on a scheme originally formulated for zooplankton as described for multiple nutrients by

Kerimoglu et al. (2020), which was based on the earlier work of Grover (2002) for a single nutrient.

Finally, respiration is calculated as a first order term, and death and lysis terms are calculated similar to those of phytoplankton:

$$R_H^{\mathrm{C}} = H^{\mathrm{C}} \zeta_H \tag{SI-28}$$

$$L_H^{\mathrm{C}} = H^{\mathrm{C}} (l_H^l + H^{\mathrm{C}} l_H^q) \tag{SI-29}$$

$$M_H^{\mathrm{C}} = H^{\mathrm{C}} (d_H^l + H^{\mathrm{C}} d_H^q) \tag{SI-30}$$

Bacteria are assumed to be immediately lysed when dead, whereas the HF and Cil are assumed to maintain their integrity, when dead. This is implemented by setting death rate of bacteria and lysis rate of the protist to 0 (Table SI-3). As in phytoplankton, lysis and death losses are channeled to $\mathrm{DOM_L}$ and $\mathrm{Det_S}$, respectively. Their corresponding N and P fluxes are calculated by multiplying the C-based rates with $Q_H^{\mathrm{N}}$ and $Q_H^{\mathrm{P}}$ (Eq. SI-2c,d, SI-4b,c).

**Table SI-3.** Parameters of the heterotroph module. Sources: A: Assumed; T: Hand-Tuned; H97: (Hansen et al., 1997); Z14: Zimmerman et al. (2014); M10:(Mahlzahn et al., 2010); dG98: Del Giorgio and Cole (1998); S: (Straile, 1997); B04: (Brown et al., 2004); O05: Oubelkheir et al. (2005).

| Symbol | Description | Value | | | | Unit | Source |
|---|---|---|---|---|---|---|---|
| | | $\mathrm{Bac_F}$ | $\mathrm{Bac_A}$ | HF | Cil | | |
| $I_{i,max}$ | Maximum ingestion rate of $\mathrm{Bac_F}$ | 4.0 | 3.0 | 4.0 | 2.4 | - | T,H97 |
| $K_i^I$ | Half saturation constant for ingestion | 10.0 | 10.0 | 20 | 20 | - | T, H97 |
| $Q_i^{\mathrm{N}}$ | Constant N:C ratio | 4.9 | 4.9 | 7.0 | 7.0 | molN $\mathrm{molC}^{-1}$ | Z14, M10 |
| $Q_i^{\mathrm{P}}$ | Constant P:C ratio | 77 | 77 | 180 | 180 | molP $\mathrm{molC}^{-1}$ | Z14, M10 |
| $\epsilon_i^{\mathrm{C}}$ | C Assimilation efficiency | 0.6 | 0.6 | 0.4 | 0.4 | - | dG98,S97 |
| $\epsilon_i^{\mathrm{N,P}}$ | N&P Assimilation efficiency | 1.0 | 1.0 | 1.0 | 1.0 | - | A |
| $d_i^l$ | Linear death rate | 0.0 | 0.0 | 0.05 | 0.05 | $\mathrm{d}^{-1}$ | T |
| $l_i^l$ | Linear lysis rate | 0.1 | 0.1 | 0.0 | 0.0 | $\mathrm{d}^{-1}$ | T |
| $l_i^q$ | Quadratic lysis rate | 0.0 | 0.0 | 0.06 | 0.02 | $\mathrm{m}^3$ (d mmolC)$^{-1}$ | T |
| $f_i^{UO}$ | Organic fraction of unassimilated excretes | 0.8 | 0.8 | 1.0 | 1.0 | - | A |
| $f_i^{UsiD}$ | Detrital fraction of unassimilated silicate | 0.9 | 0.9 | 0.95 | 0.95 | - | A |
| $\zeta_i$ | Basal respiration rate | 0.05 | | | | $\mathrm{d}^{-1}$ | A |
| $E_a$ | Activation energy for temperature scaling | 0.65 | | | | - | B04 |
| $T_{ref}$ | Reference Temperature | 283 | | | | K | - |
| $k_c^{\mathrm{H}}$ | Specific attenuation coefficient | 0.012 | | | | $\mathrm{m}^2$ $\mathrm{mmolC}^{-1}$ | O05 |

#### SI-1.1.4 Aggregation

As in Schartau et al. (2007), we consider two aggregation proccesses, denoted as $A1$ and $A2$. $A1$ represents the coalescence of $DOC_{\mathrm{S}}$ and $DOC_{\mathrm{L}}$ to form $\mathrm{TEP}^C$ (labeled with 'C' in Fig. 1). Similar to Engel et al. (2004), we consider the $\mathrm{DOC_L}$-$\mathrm{DOC_L}$ and $\mathrm{DOc_L}$-TEP, where our $\mathrm{DOC_L}$ largely corresponds to their polysaccharides (PCCHO), but in our model, $\mathrm{DOC_L}$ is a more general group that consist of hydrolyzable carbohydrates and aminoacids. In addition, we consider the interaction of $\mathrm{DOC_S}$-TEP interactions, which we assume to be less efficient than the $\mathrm{DOC_L}$-TEP interaction, while ignoring the presumably even less efficient $\mathrm{DOC_S}$-$\mathrm{DOC_S}$ and $\mathrm{DOC_S}$-$\mathrm{DOC_L}$ interactions. Inclusion of this third aggregation kernel was motivated by the observation that $\mathrm{DOC}_S$ concentrations start decreasing rapidly after an initial peak during the exponential growth phase of phytoplankton, at which the $\mathrm{Bac_A}$, the only consumers of $\mathrm{DOC_S}$, are at negligable concentrations (Fig. 2). Summing up, the first aggregation process is represented by three aggregation kernels, two of which describes the aggregation of $\mathrm{DOC_L}$ and the third that of $\mathrm{DOC_S}$:

$$A1^{\mathrm{C}}_{DOC_L} = \alpha^{A1}_{DOC_L}\beta^{A1}_{DOC_L}DOC_L^2 + \alpha^{A1}_{DOC_L-TEP}\beta^{A1}_{DOC_L-TEP}DOC_L TEP^{\mathrm{C}} \quad \text{(SI-31)}$$

$$A1^{\mathrm{C}}_{DOC_S} = \alpha^{A1}_{DOC_S-TEP}\beta^{A1}_{DOC_S-TEP}DOC_S TEP^{\mathrm{C}} \quad \text{(SI-32)}$$

here, $\beta^{A1}_{DOC_L}$ and $\alpha^{A1}_{DOC_L}$ respectively represent the collision kernel and attachment probability of interacting $\mathrm{DOC_L}$ particles, $\beta^{A1}_{DOC_L-TEP}$ and $\alpha^{A1}_{DOC_L-TEP}$ describe those of interacting $\mathrm{DOC_L}$ and TEP particles, and finally $\beta^{A1}_{DOC_S-TEP}$ and $\alpha^{A1}_{DOC_S-TEP}$ describe those of interacting $\mathrm{DOC_S}$ and TEP particles (Table SI-4). For $\beta^{A1}_{DOC_L}$ and $\beta^{A1}_{DOC_L-TEP}$, we use the values calculated by Engel et al. (2004), whereas we treat $\alpha^{A1}_{DOC_L}$, $\alpha^{A1}_{DOC_L-TEP}$, and $\alpha^{A1}_{DOC_S-TEP}$ as tuning parameters, subject to $\alpha^{A1}_{DOC_L} \leq 0.001$, $0.1 < \alpha^{A1}_{DOC_{L/S}-TEP} < 1$, following Engel et al. (2004).

$A2$ represents the aggregation of phyotplankton and detrital particles to form flocs (labelled 'F' in Fig. 1). We do not consider any biological activity within these flocs, therefore these large flocs are represented by $Det - L$. Following Ruiz et al. (2002), we describe the underlying particle dynamics with a simple, two size class model. The small particles consist of phytoplankton, TEP and small detritus, whereas the second size class consists exclusively of large detritus, i.e., flocs. Accordingly, the total aggregation rate, based on C is calculated from:

$$A_2^{\mathrm{C}} = \beta^{A2}(Phy^{\mathrm{C}} + TEP^{\mathrm{C}} + Det_S^{\mathrm{C}})Det_L^{\mathrm{C}} \quad \text{(SI-33)}$$

where $\beta^{A2}$ is the coagulation kernel, obtained as the product of a maximum value by the stickiness of the medium, determined by $TEP^C$:

$$\beta^{A2} = \beta^{A2}_{\max}\frac{TEP^{\mathrm{C}}}{TEP^{\mathrm{C}} + K^{A2}_{TEP}} \quad \text{(SI-34)}$$

Parameters $\beta_{\max}^{A2}$ and $K_{TEPC}^{A2}$ were purely empirically determined by Ruiz et al. (2002), therefore we treat them as tuning parameters (Table SI-1).

Contributions of $Phy$, $TEP$, $Det_S$ and $Det_L$ are assumed to be proportional to their concentration in the agglomerate. N, P and Si bound to the flocculating particles in $Phy$ and $Det_S$ pools are then calculated based on the stoichiometry of each of these pools:

$$A2_{Y_i}^{\mathrm{C}} = \frac{Y_i}{\sum_i Y_i} A2^{\mathrm{C}},\ Y = \{Phy^{\mathrm{C}}, TEP^{\mathrm{C}}, Det_S^{\mathrm{C}}, Det_L^{\mathrm{C}}\} \tag{SI-35}$$

$$A2_{Y_i}^{\mathrm{X}} = A2_{Y_i}^{\mathrm{C}} \frac{Y_i^{\mathrm{X}}}{Y_i^{\mathrm{C}}},\ X = \{N, P, Si\} \tag{SI-36}$$

It should be noted that the underlying coagulation process at both scales are much more complex, and depend on factors like the size distribution, and consequently the settling speed and diffusivity of interacting particles, as well as the amount of shear that is mainly determined by the turbulence in the system (Jackson, 1990). Integration of such a many-particle model is not only computationally expensive, but constraining it by observations is also difficult. Therefore we followed here the simplified two-size class approach of Ruiz et al. (2002); Engel et al. (2004); Schartau et al. (2007).

**Table SI-4.** Parameters of the abiotic material module and aggregation processes. Sources: A: Assumed; T: Hand-tuned; C: Auto-Calibrated; O05: Oubelkheir et al. (2005) for nonalgal particles; E04: Engel et al. (2004); R02: Ruiz et al. (2002).

| Symbol | Description | Value$_j$ | Unit | Source |
|---|---|---|---|---|
| $k_l$ | Specific $\mathrm{Det_L}$ leakage rate | 0.25 | $\mathrm{d}^{-1}$ | C |
| $r^{TEP}$ | Specific degradation rate of TEP | 0.1 | $\mathrm{d}^{-1}$ | A |
| $r^{Nit}$ | Specific nitrification rate | 0.2 | $\mathrm{d}^{-1}$ | T |
| $E_a$ | Activation energy for temperature scaling | 0.65 | - | B04 |
| $T_{ref}$ | Reference Temperature | 283 | K | - |
| $k_c^{\mathrm{Det}}$ | Specific attenuation coefficient | 0.012 | $\mathrm{m}^2\ \mathrm{mmolC}^{-1}$ | O05 |
| $\beta_{DOC_L}^{A1}$ | $\mathrm{DOC_L}$-$\mathrm{DOC_L}$ collision kernel for $A1$ | 0.86 | $\mathrm{m}^3\ \mathrm{mmolC}^{-1}\mathrm{d}^{-1}$ | E04 |
| $\beta_{DOC_L-TEP}^{A1}$ | $\mathrm{DOC_L}$-TEP collision kernel for $A1$ | 0.064 | $\mathrm{m}^3\ \mathrm{mmolC}^{-1}\mathrm{d}^{-1}$ | E04 |
| $\beta_{DOC_S-TEP}^{A1}$ | $\mathrm{DOC_L}$-TEP collision kernel for $A1$ | 0.032 | $\mathrm{m}^3\ \mathrm{mmolC}^{-1}\mathrm{d}^{-1}$ | A |
| $\alpha_{DOC_L}^{A1}$ | $\mathrm{DOC_L}$-$\mathrm{DOC_L}$ attachment probability for $A1$ | 0.001 | - | C |
| $\alpha_{DOC_L-TEP}^{A1}$ | $\mathrm{DOC_L}$-TEP attachment probability for $A1$ | 1.0 | - | C |
| $\alpha_{DOC_S-TEP}^{A1}$ | $\mathrm{DOC_S}$-TEP attachment probability for $A1$ | 0.85 | - | C |
| $\beta_{max}^{A2}$ | Max. collision kernel for $A2$ | 0.033 | $\mathrm{m}^3\ \mathrm{mmolC}^{-1}\mathrm{d}^{-1}$ | R02 |
| $K_{TEP}^{A2}$ | Half. sat. constant for TEP dependence of $\beta^{A2}$ | 57.48 | $\mathrm{mmolC\ m}^{-3}$ | C |

#### SI-1.1.5 Abiotic material and other general processes

Degradation of TEP to $DOM_L$ is described as a first order reaction, but without temperature dependence, as it is assumed to be a physically driven process (e.g. turbulent shear):

$$R_{TEP} = r^{TEP} TEP^{C} \tag{SI-37}$$

Nitrification of $NH_4$ into $NO_3$ is also described as a first order reaction kinetics but with temperature dependence, as it is a biologically mediated process:

$$Z = r^{Nit} \mathscr{F}_{T} NH_4^+ \tag{SI-38}$$

As the oxidation of ammonium requires oxygen, this reaction can become limited by O2 concentration in nature, however in the well mixed planktotrons, O2 never decreased below saturation levels.

All kinetic rates reported at $T_{ref}$ in Tables SI-1,SI-3 and SI-4 are scaled to the ambient temperatures in the mesocosms according to the Arrhenius equation based on the respective activation energies, $E_a$:

$$\mathscr{F}_{T} = \exp\left[\frac{E_a}{k_B}\left(\frac{1}{T} - \frac{1}{T_{ref}}\right)\right] \tag{SI-39}$$

with Boltzmann constant, $k_B = 8.6210^{-5}$ [eV K$^{-1}$].

At a given depth, $z$, the photosynthetically available radiation, is determined according to the Beer-Lambert law:

$$PAR_z = PAR_0 \exp\left(-\sum_{Y} k_c^Y Y^{C} z\right), \; Y = \{Phy, Det, Bac, Het\} \tag{SI-40}$$

where, the specific attenuation coefficient of modeled ingredients, $k_c^Y$ are listed in respective tables. $PAR_z$ is then averaged over the tank depth to calculate the $PAR$ used to calculate the light saturation of phytoplankton, $\mathscr{F}_{par}$(Eq. SI-11).

#### SI-1.2 Implementation

The model was implemented as an external library of the Framework of Aquatic Biogeochemical Models (FABM, Bruggeman and Bolding, 2014). FABM's 0D driver was used for the simulation, using measured water temperature as model forcing. For describing the light climate, $I_0$ (Eq. SI-40) was estimated using astronomical functions as described in Generalized Ocean Turbulence Model (GOTM, Burchard et al., 2006), as the light inside the mesocosms were adjusted to mimic the diurnal and seasonal variations in the field. Model was integrated using 4th order Runge-Kutta scheme with a time step of 120s.

#### SI-1.3 Model Calibration

In the following, we aim to determine a vector $\boldsymbol{\gamma}^*$ of optimal values for a number of selected parameters of the planktotron simulation model (Section SI-1). We formulate an objective function [cf. Eq. (SI-44)], which maps experimental observations, a set of parameter values $\boldsymbol{\gamma}$, and the correspondingnumerical predictions to an error value, which characterizes the goodness of the fit. The set of optimal parameters $\boldsymbol{\gamma}^*$ is then determined as a minimizer of the error function.

We define the error as a sum of component errors [cf. Eq. (SI-43)] associated to the individual observed variables. Let $\hat{\boldsymbol{x}}_X = (\hat{x}_i)_{i=1,\ldots,N_X}$ and $\boldsymbol{x}_X = (x_i)_{i=1,\ldots,N_X}$ be sequences of reference values and predicted values of the variable $X$ over the time course of the experiment, where $\hat{x}_i$, resp. $x_i$, denotes the value of $X$ at the time of the $i$-th measurment. In general, the model prediction depends on the parameters $\boldsymbol{\gamma}$ used for the numerical simulation, i.e., $\boldsymbol{x}_X = \boldsymbol{x}_X(\boldsymbol{\gamma})$.

In order to assume all variables to lie on a comparable numerical scale, we consider them to be scaled versions of the corresponding original experimental and simulated observations, $\hat{y}_{X,i}$ and $y_{X,i}$. We obtain $\hat{\boldsymbol{x}}$ and $\boldsymbol{x}$ by scaling the original variables by the standard deviation $\hat{\sigma}_X$ of the original time series. That is, $\hat{\boldsymbol{x}}_X = \hat{\boldsymbol{y}}_X/\hat{\sigma}_X$ and $\boldsymbol{x}_X = \boldsymbol{y}_X/\hat{\sigma}_X$, where,

$$\hat{\sigma}_X = \sqrt{\frac{1}{N_X}\sum_{i=1}^{N_X}\left(\hat{y}_{X,i} - \bar{y}_X\right)^2} \tag{SI-41}$$

$$\bar{y}_X = \frac{1}{N_X}\sum_{i=1}^{N_X}\hat{y}_{X,i} \tag{SI-42}$$

In our setup, the value $\hat{x}_i$ corresponds to the mean value of the observed variable $X$ at time $t_i$, where the mean is taken over the different replicates of the experiment. We define the component error function as

$$E_X(\boldsymbol{x}) = \frac{1}{N_X}\sum_{i=1}^{N_X}\frac{\left|\hat{x}_i - x_i\right|^2}{v_{X,i}} \tag{SI-43}$$

where $N_X$ is the number of observations, and $v_{X,i}$ is a normalizing factor for each point in the sequence. Note, that the function $L_X = -E_X$ can be interpreted as a likelihood if subsequent observations are assumed to be independent and the individual residuals $r_i = \hat{x}_i - x_i$ are assumed to be normally distributed with variance $v_{X,i}$, cf. Schartau et al. (2007). In order to estimate the (time-dependent) variability $v_{X,i}$, we use a combination of the variance $\hat{v}_{X,i}$ of $X(t_i)$ across replicate experiments, and an additional offset $\varrho_X > 0$. Adding this offset regularizes the residual weights, since it avoids singularities in case of $\hat{v}_{X,i} \approx 0$. Further, it represents a rough adjustment for kinematic errors originating in effects not accounted for in the model.

In order to assess parameter optimality for the model, we define the objective function $F$ as the sum of the component errors $E_X$:

$$F(\boldsymbol{\gamma}) = \sum_X w_X E_X(\boldsymbol{x}_X(\boldsymbol{\gamma})) \tag{SI-44}$$

where we allow for a different weight $w_X$ for each variable $X$, which is selected by judging the relative importance of the different components.

The variance-weighed mean of an observable,

$$\bar{x}_X = \frac{1}{N} \sum_{i=1}^{N} \frac{\hat{x}_i}{v_{X,i}} \tag{SI-45}$$

minimizes the component error $x_{\mathrm{cst}} \mapsto E_X(x_{\mathrm{cst}})$ as a function of constant model observations. Therefore, we define the total variance of variable X as:

$$V_X := E_X(\bar{x}_X) \tag{SI-46}$$

Which, in turn, can be used to determine the fraction of explained variance of the variable $X$ based on the predictors $x_i$ obtained from a parameter choice $\boldsymbol{\gamma}$ as:

$$R_X^2(\boldsymbol{\gamma}) = 1 - \frac{E_X(\boldsymbol{x}(\boldsymbol{\gamma}))}{V_X} \tag{SI-47}$$

Then, the overall fraction of explained variance of the full observation set can be calculated as:

$$R^2(\boldsymbol{\gamma}) = 1 - \frac{F(\boldsymbol{\gamma})}{V}, \tag{SI-48}$$

where $V = \sum_X w_X V_X$.

For obtaining the optimal fit, we considered the errors of all variables excluding Total DOC, Chl $a$,

DON, NH4, for which weights, $w$ were assumed to be 0.0 (Table SI-6). Exclusion of Total DOC was to avoid a double counting of the DOC, given that the $\mathrm{DOC_S}$ and $\mathrm{DOC_L}$ sum up to the concentration of Total DOC. The reason for excluding Chl-$a$ was the observation that the Chl-$a$ concentrations during the decline phase were maintained at relatively high concentrations in comparison to the POC, presumably caused by pigments in the phytodetrital aggregates being preserved fo a while (e.g. Passow et al., 2003). In our model, the pigments in the flocculated phytoplankton is assumed to be degraded immediately, as describing the Chl $a$ within $\mathrm{Det_L}$ and its degradation would require an additional state variable and parameter(s). Finally, DON and NH4 were excluded because it appears as if capturing the low concentrations of these two variables (and that of DIP) towards the end of the experiment would require consideration of flexible stoichiometry of bacteria (Godwin and Cotner, 2015). The inability of the model to reproduce the course of these three variables is therefore to be expected. Their inclusion in the calibration results in lower amount of errors in these variables but likely for wrong reasons, and at the cost of higher errors in other variables. Therefore we decided to disregard these variables.

We perform the minimization of (SI-44) by manually providing an initial parameter guess first to the Powell method (Powell, 1964) and then the Melder-Nead method (Gao and Han, 2012), both implemented in SciPy 1.8.0. We simultaneously optimized eight parameters as listed in Table SI-5,chosen based on ($i$) a high degree of uncertainty of their values, which are presumably strongly dependent on the specific experimental conditions, such that they cannot be adopted from the literature; ($ii$) a preliminary sensivity analysis that demonstrated the sensitivity of the results to these parameters. For the optimization postulated plausible ranges, mainly from literature research. We implemented these ranges as hard bounds to the optimization, that is only values inside the given interval are admissible but no penalties were imposed towards the boundary.

**Table SI-5.** Optimal fits and employed admissible ranges for parameter calibration.

| Symbol | Description | Unit | Optimal fit $\gamma_j^*$ | Range |
|---|---|---|---|---|
| $l_P^l$ | Specific lysis rate | $d^{-1}$ | 0.1 | [0.1, 0.3] |
| $\gamma_{pp}$ | Production sp. respiration rate | - | 0.11 | [0.0, 0.2] |
| $K_{TEP}^{A2}$ | Half sat. constant for TEP dependence of $\beta^{A2}$ | mmolC $m^{-3}$ | 57.48 | [10, 100] |
| $\alpha_{DOC_L}^{A1}$ | $\mathrm{DOC_L}$-$\mathrm{DOC_L}$ attachment probability for A1 | - | 0.001 | [0, 0.001] |
| $\alpha_{DOC_L-TEP}^{A1}$ | $\mathrm{DOC_L}$-TEP attachment probability for A1 | - | 1.0 | [0.1, 1] |
| $\alpha_{DOC_S-TEP}^{A1}$ | $\mathrm{DOC_S}$-TEP attachment probability for A1 | - | 0.85 | [0.0, 1] |
| $k_l$ | Specific $\mathrm{Det_L}$ leakage rate | $d^{-1}$ | 0.25 | [0.0, 0.25] |
| $\mathrm{TEP}_0^C$ | Initial $\mathrm{TEP^C}$ concentration | mmolC $m^{-3}$ | 0.62 | [0.0, 2.0] |

For the optimal fit, weighed sum of the component errors, $F(\boldsymbol{\gamma}) = 0.786$, and the overall fraction of explained variance is $R^2 = 0.905$ for the calibration variables displayed in Fig. 2 (See Fig. SI-2 for a comparison of observed and simulated variables that were excluded from model calibration).

Characteristics of the optimal fit for individual observables are reported in Table SI-6.

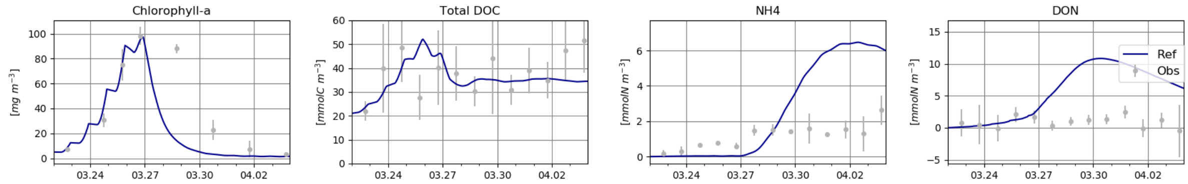


**Figure SI-2.** Observed and simulated variables that were ignored in model calibration.

**Table SI-6.** For the optimal fit, characteristics for individual observables used in the calculation of the error function (Eq. SI-44).

| **Variable** | weight $w_X$ | avg. cross-replicate variance $\langle \hat{v}_{X,i} \rangle$ | component error $E_X^*$ | $R_X^2$ |
|---|---|---|---|---|
| NO3 | 1.0 | 0.205122 | 0.038 | 0.961 |
| DIP | 1.0 | 0.147233 | 0.060 | 0.936 |
| DISi | 1.0 | 0.0326602 | 0.003 | 0.997 |
| $DOC_S$ | 1.0 | 1.00433 | 0.082 | 0.811 |
| $DOC_L$ | 1.0 | 0.720968 | 0.067 | 0.887 |
| $Bac_F$ | 1.0 | 0.469645 | 0.070 | 0.881 |
| HF | 1.0 | 0.374799 | 0.121 | 0.821 |
| Cil | 1.0 | 0.616351 | 0.108 | 0.57 |
| POC | 1.0 | 0.228494 | 0.124 | 0.870 |
| PON | 1.0 | 0.261417 | 0.033 | 0.966 |
| POP | 1.0 | 0.295917 | 0.081 | 0.913 |
| Total DOC | 0.0 | 1.4063 | 0.245 | 0.413 |
| Chl-$a$ | 0.0 | 0.157619 | 0.448 | 0.538 |
| NH4 | 0.0 | 0.583769 | 6.680 | -9.599 |
| DON | 0.0 | 2.02086 | 12.168 | -48.731 |

### SI-2 Supplemental Figures

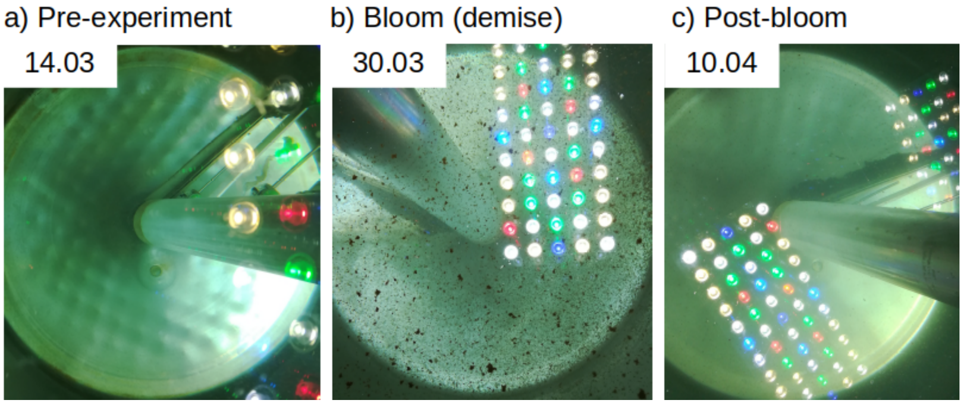


**Figure SI-3.** Photographs of tanks taken from the surface throughout the experiment.

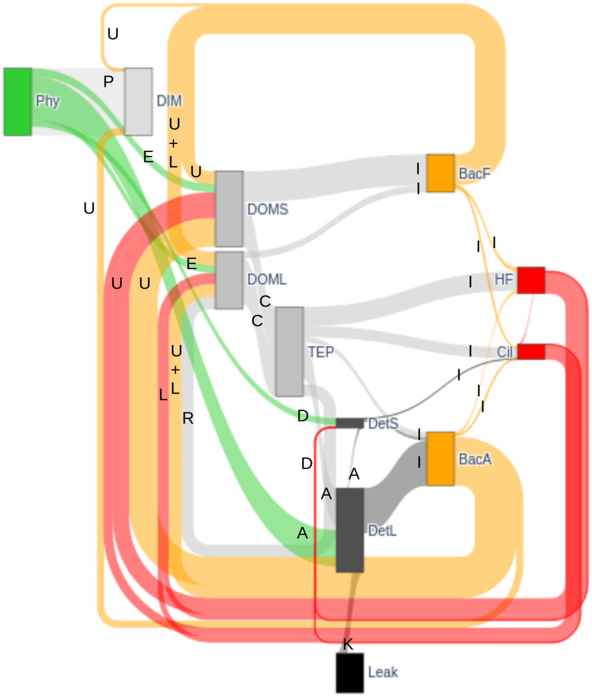


**Figure SI-4.** Lava plots (a.k.a. Sankey diagrams) of simulated fluxes, where the width of links between model components are scaled with the total flux between the components throughout the simulation. Abbreviations/annotations as in Fig. 1.

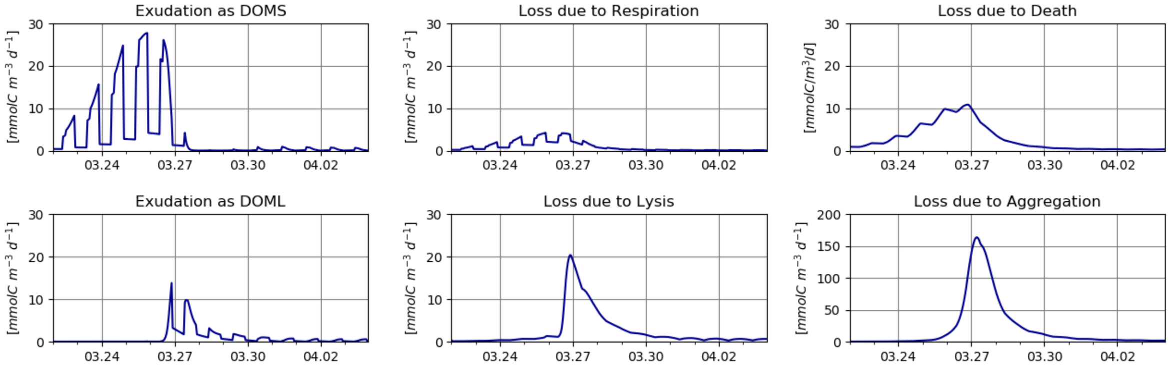


**Figure SI-5.** Diatom loss rates as estimated by the model.

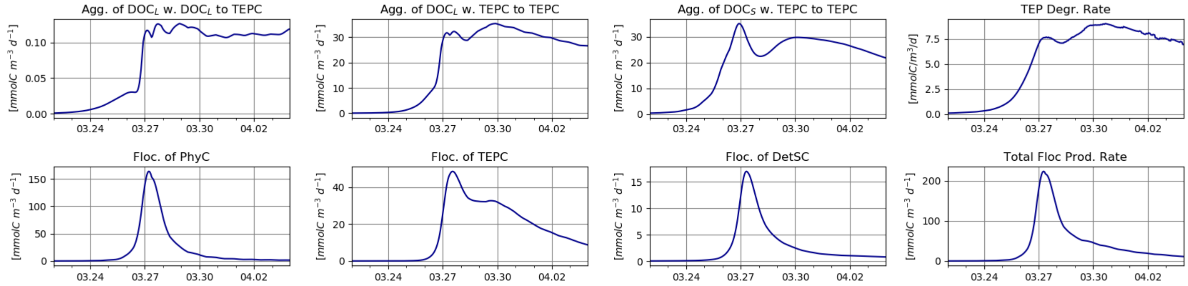


**Figure SI-6.** Dynamics of TEP (top row) and floc (bottom row) formation

.

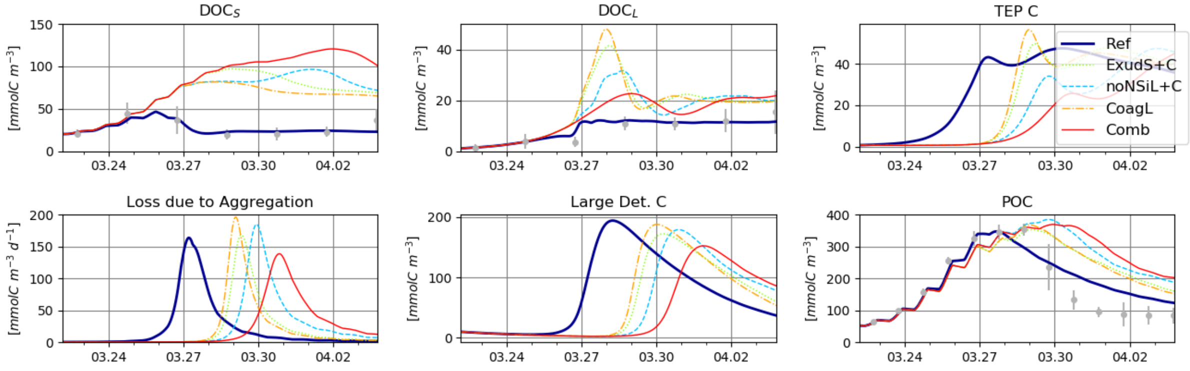


**Figure SI-7.** As in Fig. 5, but each of 'ExudS' and 'NSiL' scenarios combined with 'CoagL' scenario. 'Ref', 'CoagL' and 'Comb' runs are identical to those in Fig. 5, and shown here again only for convenience.

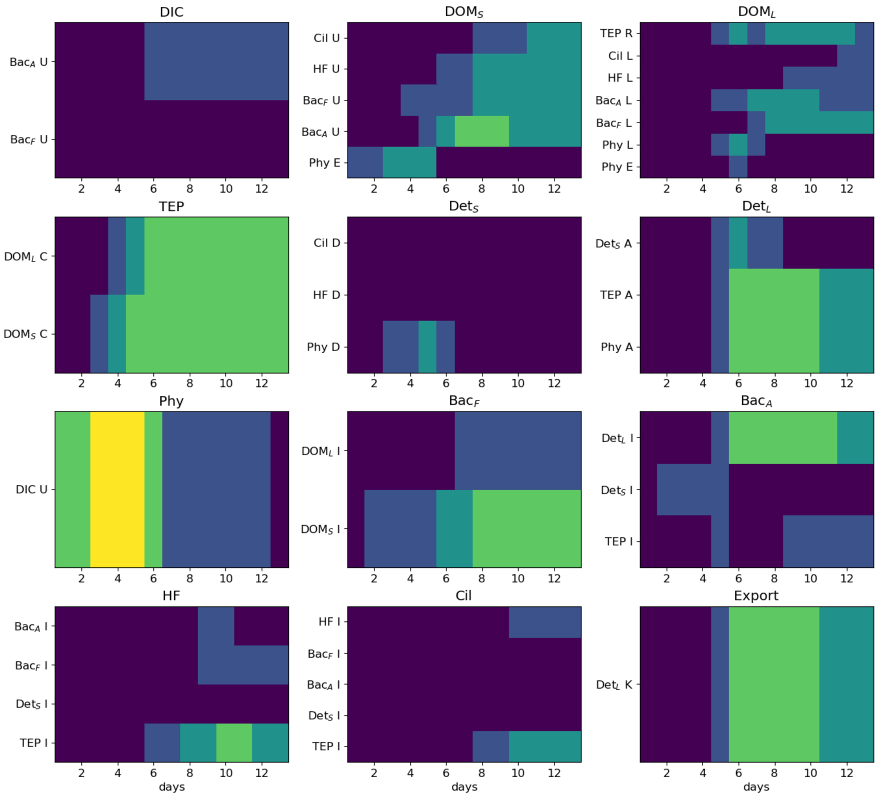


**Figure SI-8.** Temporal course of individual influxes driven by each source and process (abbreviations as in Fig. 1), as simulated by the reference run.
